# Abnormal homeostasis of P53 gene knockout mice can be reflected in urinary proteome

**DOI:** 10.1101/2023.04.29.538827

**Authors:** Xuanzhen Pan, Minhui Yang, Yijin Bao, Yongtao Liu, Youhe Gao

## Abstract

In this experiment, the urinary proteome analysis of p53 knockout mice at four relatively early time points before death was performed from three dimensions. One was to compare the differential proteins screened by homozygote and heterozygote at the same time point; the other was to compare the differential proteins screened by homozygote and wild-type as well as heterozygote and wild-type at the same time point; the third was to compare the differential proteins screened by homozygote and heterozygote before and after each time point, and to screen the differential proteins of three mice by self-control analysis and screening. Later, enrichment analysis of the differential proteins was performed mainly by Metascape and String, and disease association analysis of gene or variant genotype-phenotype with the help of DisGeNET and Monarch. We found that the biological pathways mainly related to metabolism were most abundant, and at the same time abnormal homeostasis, cancer and tumor, cardiovascular diseases, and neurodegenerative diseases appeared. However, under the analysis of Monarch, homozygotes or heterozygotes contained abnormal homeostasis at all time points, and most of them ranked the first or at least the top, while all timepoints in the growth and development of each normal rat were not enriched to abnormal homeostasis and other disease pathways.

## Introduction

TP53 is a key tumor suppressor gene that mutates in more than half of human cancers [1, 2].

The mutation of TP53 not only weakens its antitumor activity, but also confers carcinogenic properties to the mutant p53 protein. The function of p53 in normal cells. P53 is an important tumor suppressor for maintaining homeostasis in normal cells. Throughout its life cycle, cells are subjected to sustained stress, both endogenous and exogenous. To overcome these stresses, p53 is activated to mediate a range of cellular responses via its transcription-dependent function or direct protein-protein interaction [3]. The gene plays an irreplaceable role in cell cycle, autophagy, apoptosis, redox homeostasis, aging, metabolism, and other aspects, as shown in Figure 1 [4]. Many endogenous and exogenous stressors can activate p53, triggering its further regulation of a series of cellular responses necessary to maintain homeostasis in the body. P53 knockout mice can trigger a series of cancer tumors, such as fibrosarcoma, thymic lymphoma, pancreatic cancer, lung cancer, etc. [5–8]. Here, we collected the urine of homozygous and heterozygous p53 knockout mice and the urine of wild-type mice for comparative analysis.

**Fig. 1.**
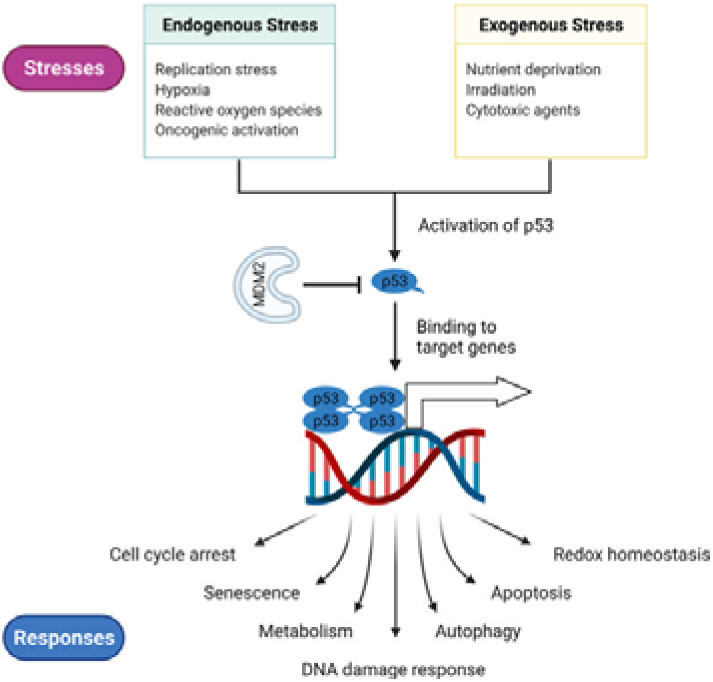
The functions of p53 in the normal cells.

## Materials and methods

### Acquisition of experimental sample

Two pairs of heterozygotes of C57BL/6J-Trp53 living mice (strain name: KOCMP-20952-Trp53) were purchased from Santa Clara. and then entrusted to Beijing Charles River Laboratory. to carry out surrogate rearing and subsequent breeding identification with the two pairs of mice as the initial female parent. The parent animals were raised by permanent cohabitation, and the neonatal rats were divided into cages according to gender after their milk was cut off. After the parents retire, the appropriate provenance of the next generation will be selected as the breeding parent, and two generations of animals will be preserved simultaneously during conservation. The animal experiments were approved by the Ethics Review Committee of the Institute of College of Life Science, Beijing Normal University, China. All experimental animals were utilized following the “Guidelines for the Care and Use of Laboratory Animals” issued by the Beijing Office of Laboratory Animal Management (Animal Welfare Assurance Number: ACUC-A02-2015-004).

These urine samples were collected in the same environment, frozen in −80°C fridge refrigerators, and then processed together.

### Identification of Homozygote, Heterozygote and Wild Type

Primers sequence:

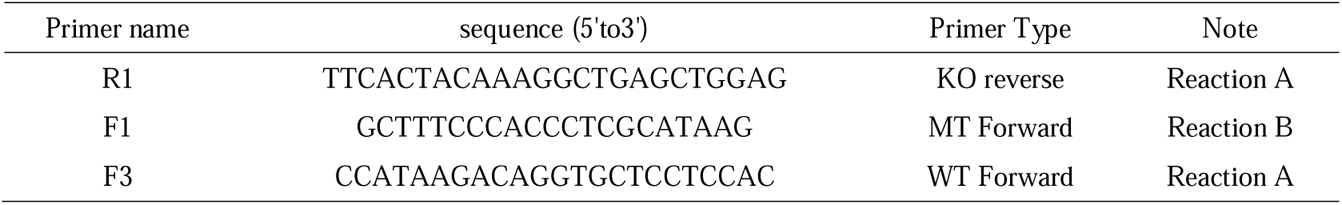

Criteria for judgment:

Homozygote: 1063 bp; Heterozygote: 1063 bp/759 bp; Wild Type: 759 bp.

PCR condition and system:

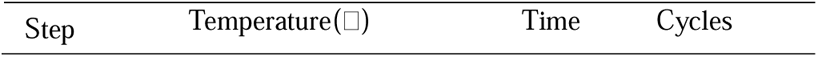
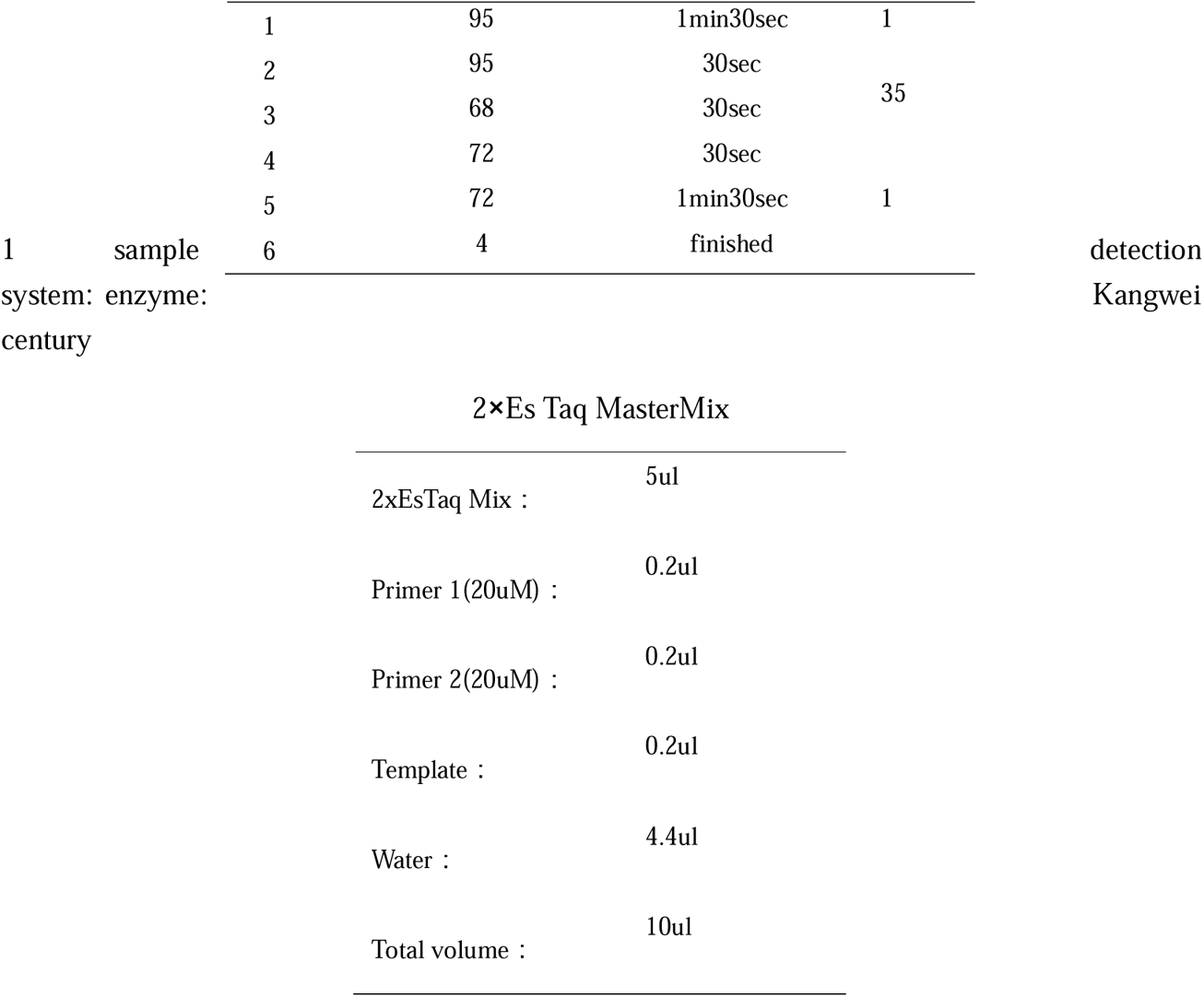

The PCR results show that:

As shown in Figure 2 below, No.6 is the wild type, No.7 and No.9 are homozygotes, and others are heterozygotes.

**Fig. 2.**
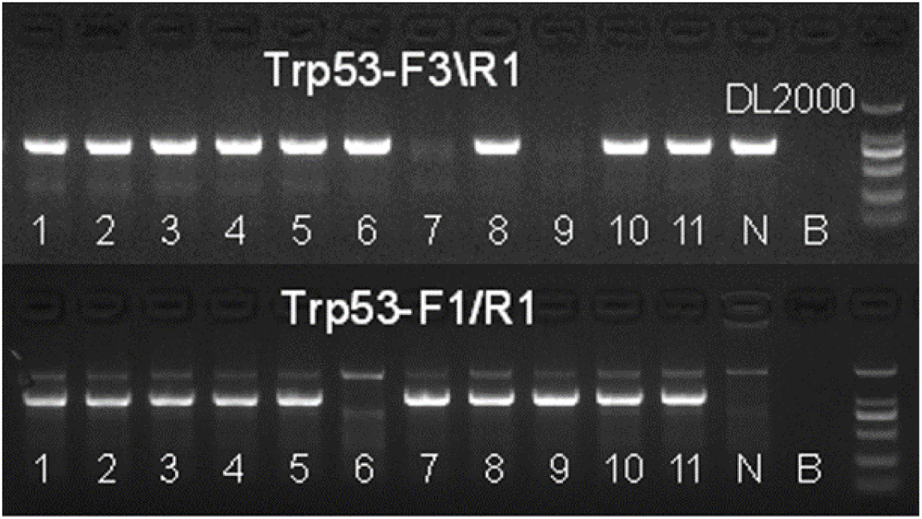
Pcr gel electrophoresis showed homozygosity and so on.

Using the instruments of Institute of Animal Science of Chinese Academy of Sciences, CT (computed tomography) was performed on some small animals and the films of various parts of the whole body of five mice 16, 24, 28, 40 and 41 were taken. It was found that only mouse No.40 could see a clear tumor mass on the upper side of the kidney, and mouse No.40 was homozygous, as shown in the right figure of Figure 3 below. The films of the scans of other mice from the brain to the whole body showed no visible tumor formation, as shown in the left figure of Figure 3 below.

**Fig. 3.**
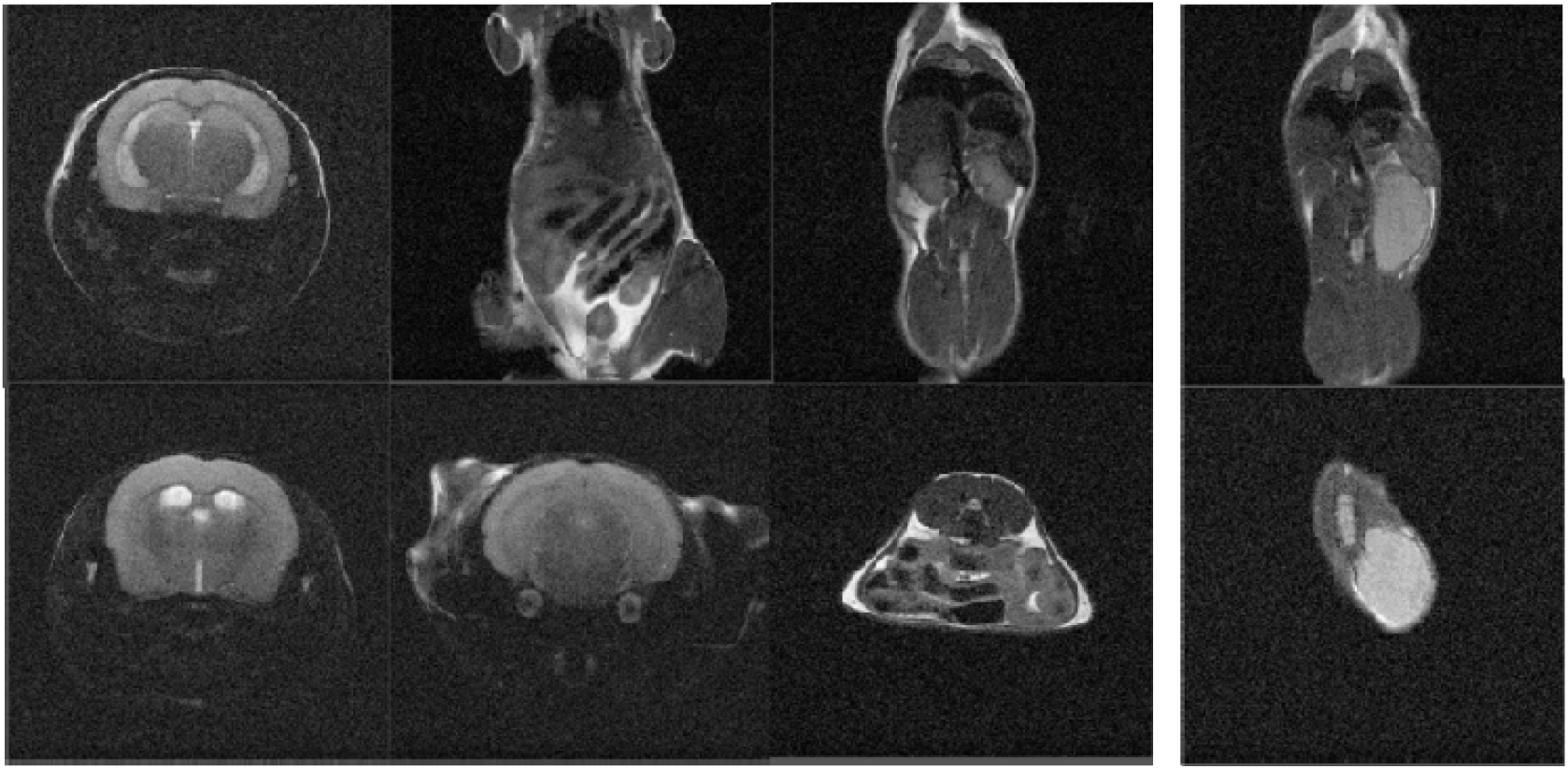
The CT scans of the whole body of the mouse are normal on the left, and the tumor on the right.

### Preparation of protein sample and trypsin digestion

Urine samples from gene-knockout mice were reacted with 20 mmol/L dithiothreitol (DTT) at 37 C for 1 h to denature disulfide bonds in the protein structure, followed by the addition of 55 mmol/L iodoacetamide (IAA) and reaction in the dark for 30 min to alkylate disulfide bond binding sites. The supernatant was precipitated at-20 C with three volumes of pre-cooled acetone for 2 h and then centrifuged at 4 C and 12,000 ×g for 30 min to obtain a protein precipitate. The precipitate was then resuspended in a suitable amount of proteolysis solution (8 mol/L urea, 2 mol/L thiourea, 25 mmol/L DTT and 50 mmol/L Tris). The concentration of that protein extract was measure using Bradford analysis. By using method 15 of filter-assisted sample preparation (FASP), each sample was hydrolyzed to 100 µg of protein in a 50: 1 ratio using trypsin (trypsin Gold, MassSpec Grade, Promega, Fitchburg, WI, USA). After enzymolysis at 37 for 14 h, 10% formic acid solution was added to the solution to terminate the enzymolysis, and the polypeptide solution was obtained after centrifugation through a 10 KDa ultrafiltration tube. The concentration of the polypeptide was determined by using the BCA method, and dried by a vacuum centrifuge concentrator (Thermo Fisher, USA). The dried polypeptide was sealed and stored at −80□.

### Liquid chromatography and Mass spectrometry

Before analysis of urine samples of gene-knockout mice, the dried peptide samples should be dissolved in 0.1% FA(formic acid) for liquid chromatography-mass spectrometry analysis, the final concentration should be controlled at 0.1 μg/μL, and each sample should be analyzed with 1 μg peptide. For DDA (data-dependent acquisition) experiments, iRT (indexed retention time; Biogenesis, Switzerland) calibration peptides were spiked into the sample. Thermo EASY-nLC1200 chromatography system was loaded to Pre-column and the analytical column. Proteome data was collected by the Thermo Orbitrap Fusion Lumos mass spectrometry system (Thermo Fisher Scientific, Bremen, Germany). Liquid chromatography analysis method: pre-column: 75 μm×2 cm, nanoViper C18, 2 μm, 100Å; analytical column: 50 μm×15 cm, nanoViper C18, 2 μm, 100 Å; injection volume: 10 μL, flow rate: 250 nL/min. The mobile phase configuration is as follows, phase A: 100% mass spectrometric grade water (Fisher Scientific, Spain)/1‰ formic acid (Fisher Scientific), phase B: 80% acetonitrile (Fisher Scientific, USA)/20% water/1‰ formic acid, 120 Min gradient elution: 0 min, 3% phase B; 0 min-3 min, 8% phase B; 3 min-93 min, 22% phase B; 93 min-113 min, 35% phase B; 113 min-120 min, 90% phase B; mass spectrometry method, ion source: nanoESI, spray voltage: 2.0 kV, capillary temperature:

320°C, S-lens RF Level: 30, resolution setting: level 1 (Orbitrap) 120,000 @m/z 200, Level 2

30,000 (Orbitrap) @m/z 200, precursor ion scan range: m/z 350-1350; product ion scan range: from m/z 110, MS1 AGC: 4e5, charge range: 2-7, Ion implantation time: 50 ms, MS2 AGC: 1e5, ion implantation time: 50ms, ion screening window: 2.0 m/z, fragmentation mode: high energy collision dissociation (HCD), energy: NCE 32, Data-dependent MS/MS : Top 20, dynamic exclusion time: 15s, internal calibration mass: 445.12003.

### Mass spectrometry data processing

The Swiss-Port protein database of Musculus (mouse/house mouse/laboratory mouse) downloaded from Uniprot website was imported into Spectronaut to construct the database. The RAW data of DIA-MS was imported into Spectronaut software for analysis, and the default parameters of the software were used. The software calculates the retention time of peptide fragments according to iRT data. By comparing other parameters such as retention time and mass-to-core ratio with the data in the spectrum library, the Q value was set to 0.01 threshold, which corresponds to FDR<0.01. According to the protein of the specific peptide fragments filtered by Q value, the peptide fragment intensity was calculated by the peak area of each fragment ion of MS2, and the corresponding protein intensity was obtained by integrating and summing the peptide fragment intensities to achieve the quantitative purpose. The software would automatically normalize the quantitative preliminary data. The quantitative protein intensity level distribution was shown in Figure 5 below, which spaned at least 6 orders of magnitude after taking the logarithm of 10. Screening differential protein, introducing the differential protein into online website such as DAVID(https://david.ncifcrf.gov), Metascape(http://metascape.org/), and others performed biological pathway analysis and functional annotation evaluation, including annotation of protein’s molecular functions, cellular components, and biological processes. An analysis of the interaction between protein was performed using the online site string (https://string-db.org). Because the sample includes four earlier time points (22 days T1, 42 days T2, 62 days T3 and 70 days T4 after weaning), homozygotes, heterozygotes and wild types exist at the same time, so we try to analyze and compare them comprehensively from three angles. Firstly, the comparative analysis of homozygote and heterozygote samples at the same time point; Then the homozygotes and heterozygotes were compared with the wild type at each time point. Finally, homozygote and wild type were analyzed according to the comparison before and after time.

**Fig. 4.**
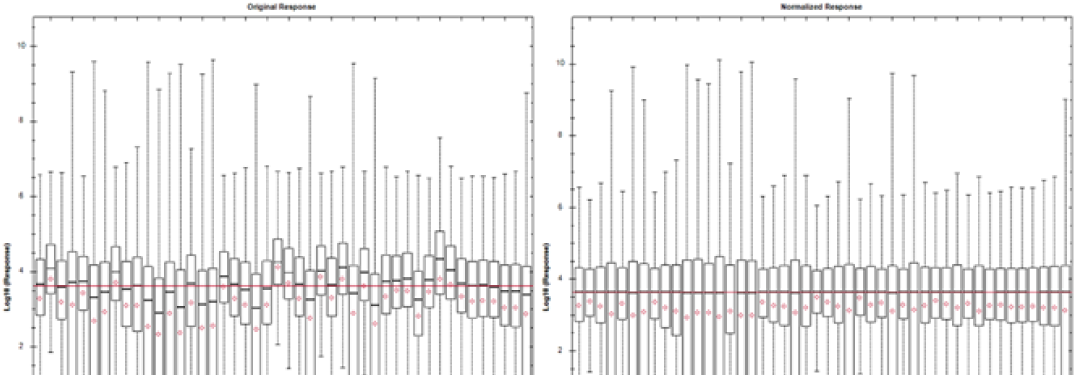
Normalization diagram of raw data

**Fig. 5.**
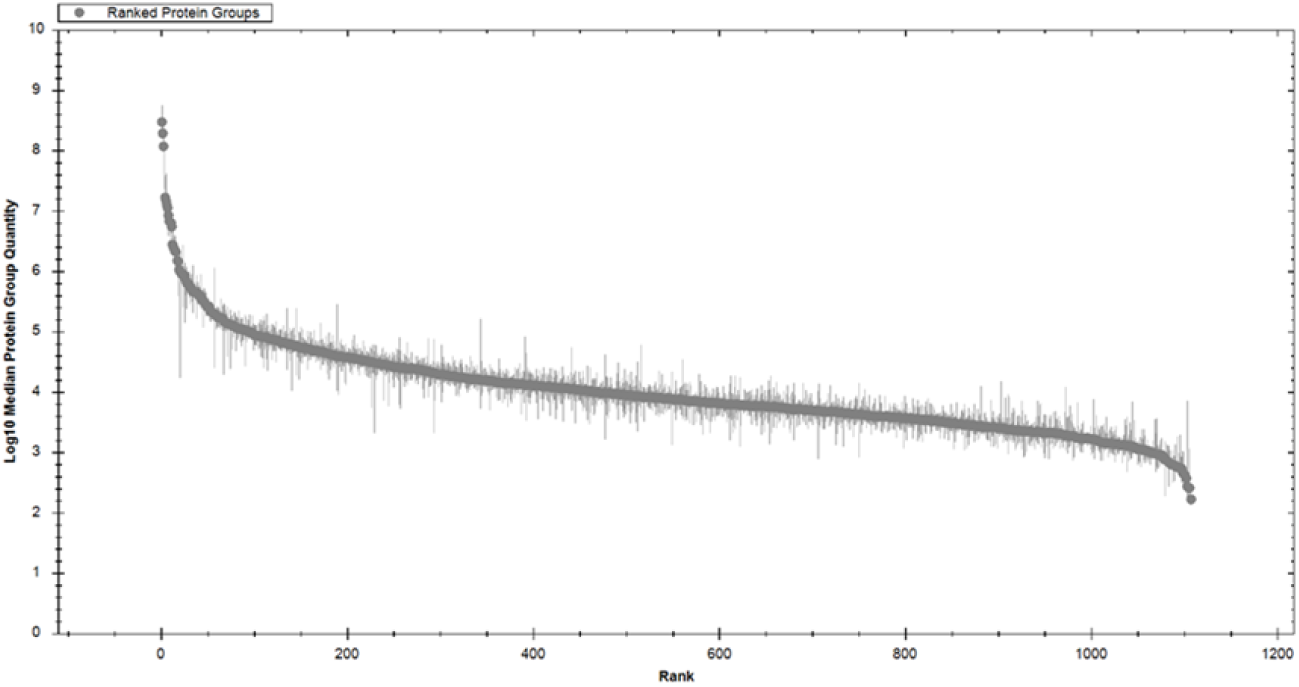
The magnitude span of the quantified protein intensities

## Results

### Comparative analysis of homozygotes and heterozygotes at the same time point

Separate analysis was performed for each time point first. The homozygotes and heterozygotes in each time point were grouped and compared. Screening was performed based on the p value < 0.05 and fold-change value < 0.5 or > 2 obtained from the two-tailed t-test, to obtain eligible differential proteins and the protein unique to homozygotes and heterozygotes. Homozygote and heterozygote were compared at each time point to obtain differential proteins. The intersection of the differential proteins at the four time points is shown in Fig. 6. There were only two proteins in common at the four time points, and there were more unique differential proteins at the four time points, which also revealed the difference complexity between homozygote and heterozygote after p53 gene knockout.

**Fig. 6.**
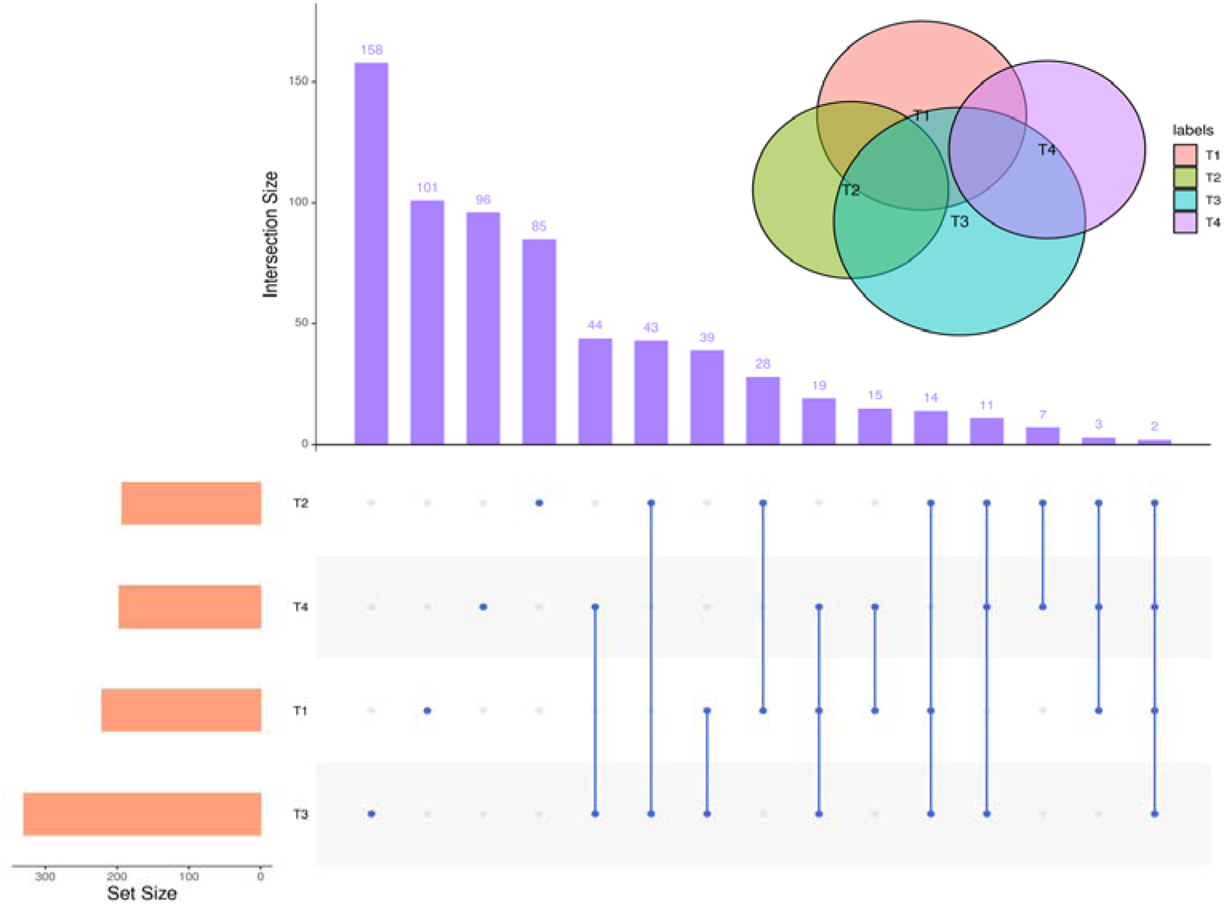
The interaction between homozygote and heterozygote differential proteins at four time points

At the first time point, 22 days after weaning, we imported the obtained differential proteins to the Metascape website for enrichment analysis using the default parameters. The following Table 1 lists the databases used in the latest version of Metascape to retrieve their annotations [9].

**Table 1.** Gene annotations extracted

| Table. 1 Gene annotations extracted |  |  |
| --- | --- | --- |
| Name | Type | Description |
| Gene Symbol | Description | Primary HUGO gene symbol. |
| Description | Description | Short description. |
| Biological Process (GO) | Function/Location | Descriptions summarized based on gene ontology database, where up to three most informative GO terms are kept. |
| Kinase Class (UniProt) | Function/Location | Detailed kinase classes. |
| Protein Function (Protein Atlas) | Function/Location | Protein Function (Protein Atlas) |
| Subcellular Location (Protein Atlas) | Function/Location | Subcellular Location (Protein Atlas) |
| Drug (DrugBank) | Genotype/Phenotype/Disease | Drug information for the given gene as target. |
| Canonical Pathways | Ontology | Canonical Pathways |
| Hallmark Gene Sets | Ontology | Hallmark Gene Sets |

Enrichment analyses for each given gene list, pathway, and biological process were performed using the following ontological sources: Kegg Paths, Go Biological Procedures, Reach ome Gene Sets, Canonical Paths, Corum, Wiki Paths, and Panther Paths. All differential proteins entered were used as background for enrichment analysis. The terms with p value < 0.01, minimum count of 3, and enrichment factor > 1.5 (the ratio between observed and occasionally expected counts) were restricted and clustered into groups based on their member similarities. More specifically, the p-value was calculated based on the cumulative hypergeometric distribution, and the q-value was obtained by multiple comparisons using the Benjamini-Hochberg method. When hierarchical clustering was performed on the enrichment terms, the Kappa coefficient was used as the similarity measure, and a subtree with a similarity > 0.3 was considered as a branch cluster. The most statistically significant item in each cluster was selected to represent the cluster [10, 11]. As shown in Fig. 7 below, representative pathways for enrichment analysis in the first 20 branches obtained by using the appeal idea are shown, while Fig. 8 shows the network relationship among these representative pathways, where pathways with similarity > 0.3 are connected, with no more than items shown in each cluster. The network is visualized using Cytoscape [12]. The colors of the representative pathways under each cluster classification are different as the left figure, and they are ordered from shallow to deep as the right figure according to the p value of each pathway.

**Fig. 7.**
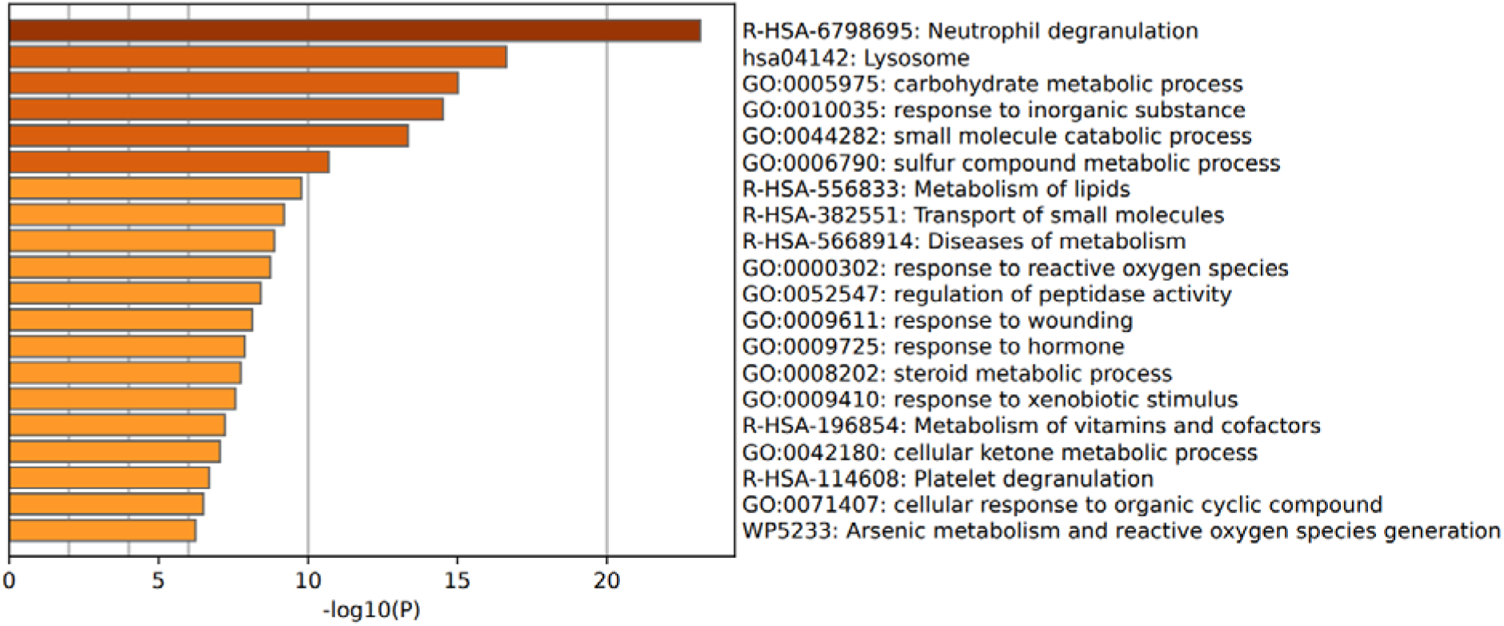
Top 20 clusters with their representative enriched terms (one per cluster, T1 Homozygote-Heterozygote)

**Fig. 8.**
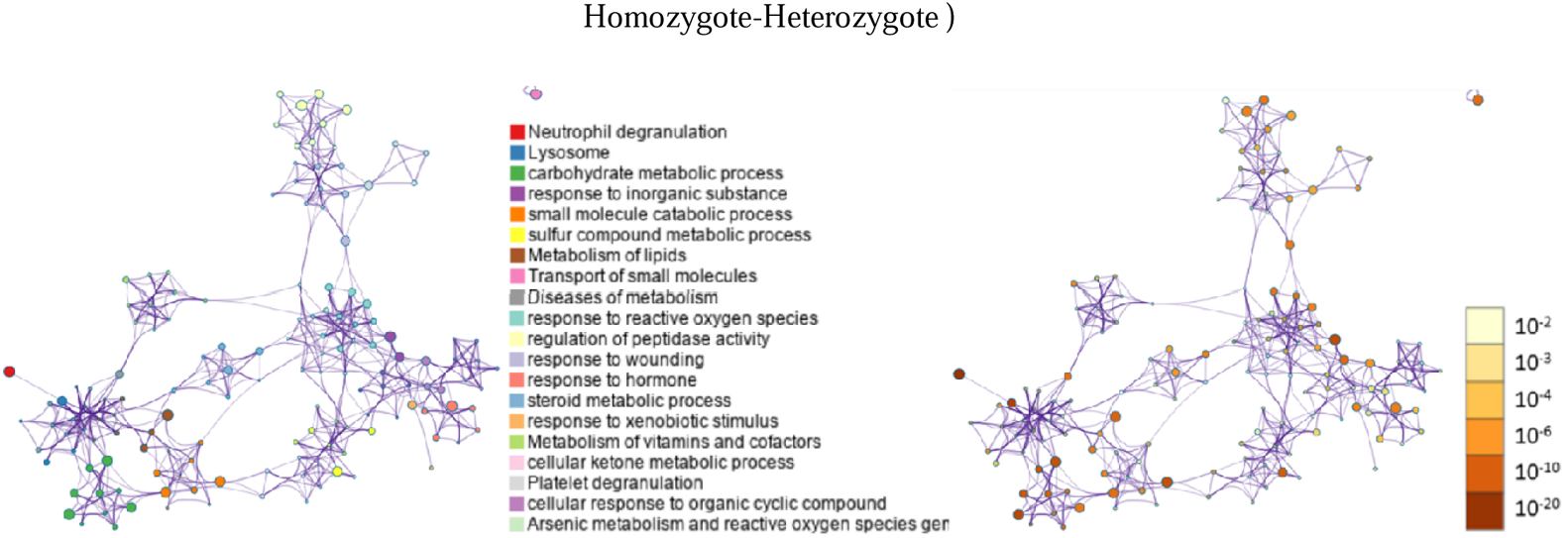
Network of enriched terms: (left) colored by cluster ID, where nodes that share the same cluster ID are typically close to each other; (right) colored by p-value, where terms containing more genes tend to have a more significant p-value. (T1, Homozygote-Heterozygote)

As shown in Fig. 8, neutrophil degranulation process was ranked first with the lowest P value, while GO biological process focused on metabolic processes, including carbohydrate metabolism, small molecule metabolism and sulfur-containing compound metabolism. Next, the pathways of Reactome also involve in lipid metabolism, small molecule transport, and metabolic diseases. Finally, there are other biological processes such as steroid metabolism, vitamin and coenzyme factor metabolism, and cellular ketone metabolism. It can be seen that eight analyses in the first 20 years were metabolic, while the analysis of Reactome directly mentioned metabolic diseases. The known studies also show that p53 has the function of metabolic regulation [13], and metabolic changes are considered to be the marker of tumors and the key factors for the occurrence and development of tumors. It is known that P53 regulates metabolism in many different ways, including glycolysis, lipid metabolism, mitochondrial oxidative phosphorylation, pentose phosphate pathway, fatty acid synthesis and oxidation [14], in order to maintain the steady state of cell metabolism. However, knocking out the urine protein of mice by p53 may provide a new way to explore other metabolic functions of p53 gene. For example, our preliminary exploration has revealed that p53 may affect other metabolism such as steroids and ketones. As shown in Figure 9 below, biological pathways with a top P value < 0.01 in GO analysis were displayed. Metabolic processes were ranked first, followed by steady-state processes, as well as immune system processes and growth and development. Therefore, after p53 knock-out, the urine protein group of mice showed such a strong effect of gene knock-out that it destroyed the body’s steady state and metabolic processes, which were consistent with the findings of existing studies.

**Fig. 9.**
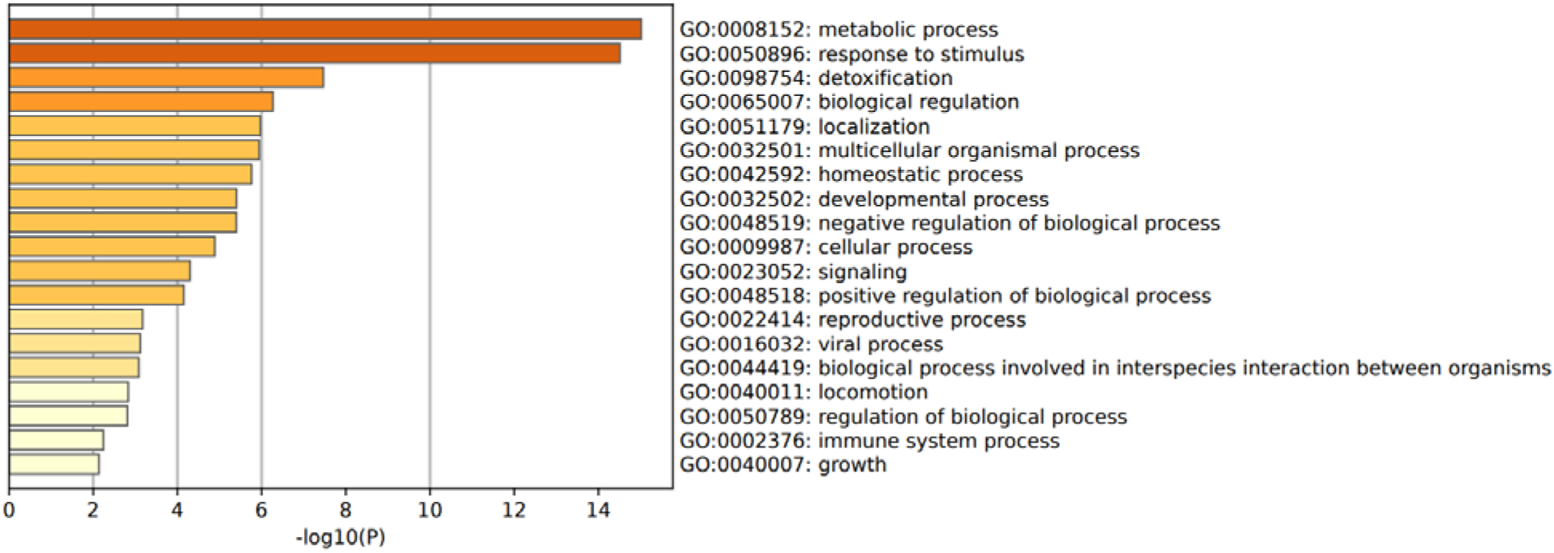
The top-level Gene Ontology biological processes (T1, Homozygote-Heterozygote)

Through DisGeNET [15], differential proteins are analyzed for gene and disease association, and as shown in Fig. 10 below, a series of top-ranking related diseases are shown, including fatty liver, lysosomal storage disease, diarrhea, hepatomegaly, malnutrition, pre-senile dementia, myocardial ischemia, drug-induced liver disease, malaria, acute kidney injury, motor neuron disease, enzyme disease, delayed Alzheimer’s disease, thoracic protuberant tongue, acute renal insufficiency, vomiting, diabetes, endothelial dysfunction, and Pooch’s syndrome, i.e., multiple bone dysplasia syndrome and hemolytic anemia. It could be seen that only in the short term after lactation, the deletion of p53 gene caused great damage to the body, and the homozygosity was more serious than heterozygosity.

**Fig. 10.**
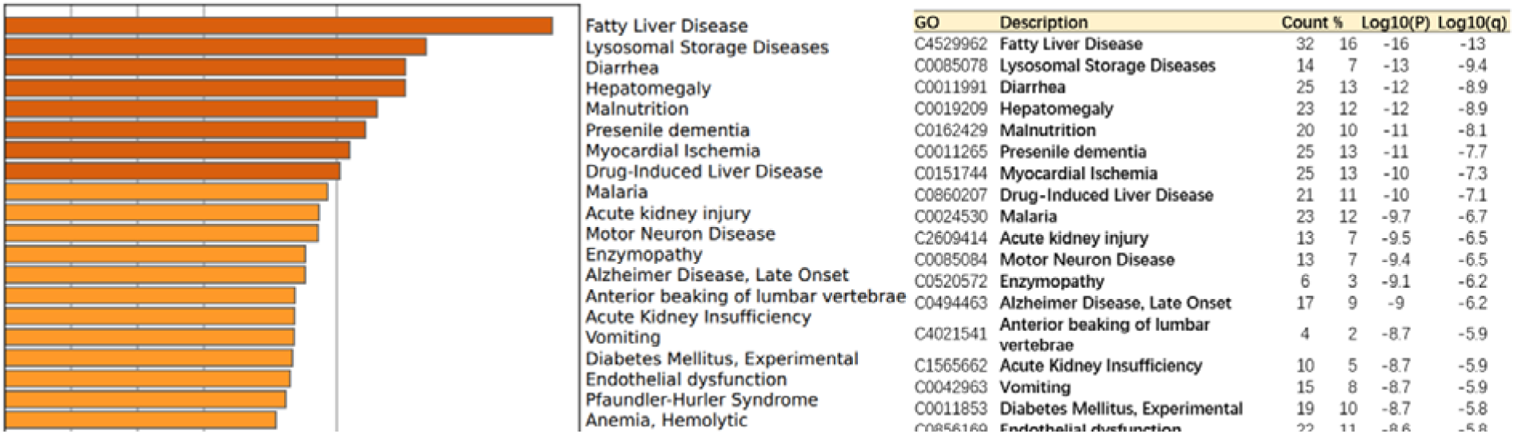
Summary of related disease enrichment analysis in DisGeNET (T1, Homozygote-Heterozygote)

Next, the differential proteins at each time point of T2, T3 and T4 were introduced into the Metascape website for enrichment function analysis, as shown in Fig. 11. We found that they were enriched to steady-state processes, including multi-cell steady-state and bone marrow cell steady-state, and there were also stress-and coagulation-related pathways. In time point of T4, there were more metabolic processes, such as sulfur-containing compound metabolism, and small-molecule metabolism, including nucleobase-containing small-molecule metabolism, monocarboxylic acid metabolism, alcohol metabolism, glycosyl compound metabolism, and glycolipid catabolism. Disease-related analysis at the four time points was enriched to a total of 69 credible disease types, as shown in Table 2 below, with the count value being the number of input differential proteins in the disease. Amyloidosis was abundant in the latter three time points, followed by the diseases existing in the two time points such as lysosomal storage disease, senile dementia, drug-induced liver disease, endothelial dysfunction, vascular disease, lupus nephritis, dermatosis, atopic dermatitis, skin damage, mesothelioma and Behcet’s syndrome. Then there were the diseases that were enriched individually at each time point, involving myocardial infarction, thrombosis, cystemia, meningioma, lymphoma, renal cell carcinoma, prostate carcinoma, hepatitis, thyroid carcinoma, malignant mesothelial carcinoma, aortic aneurysm, colonic adenocarcinoma, etc.

**Fig. 11.**
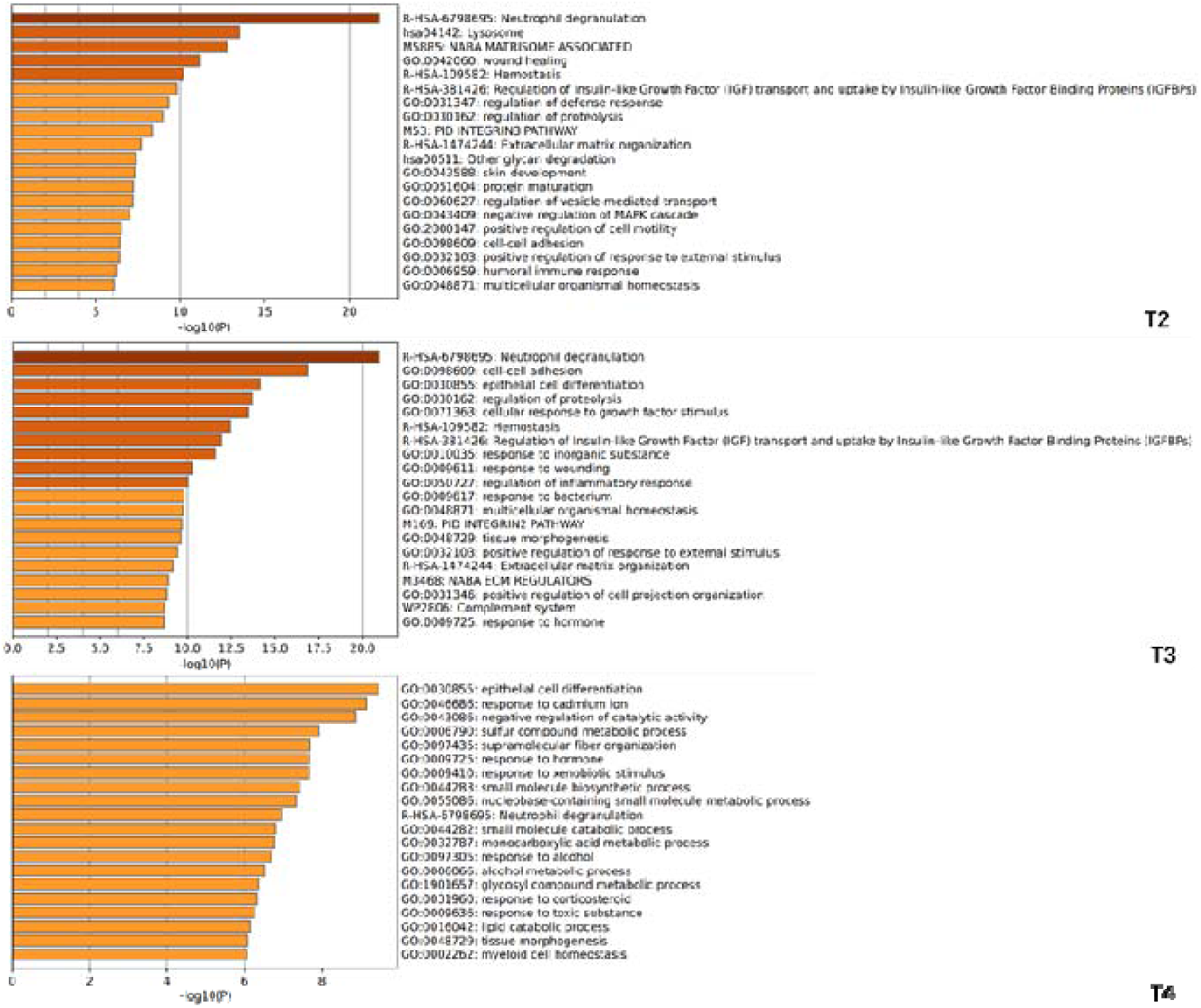
Top 20 clusters with their representative enriched terms (one per cluster, T2~T4 Homozygote-Heterozygote)

**Table 2.**
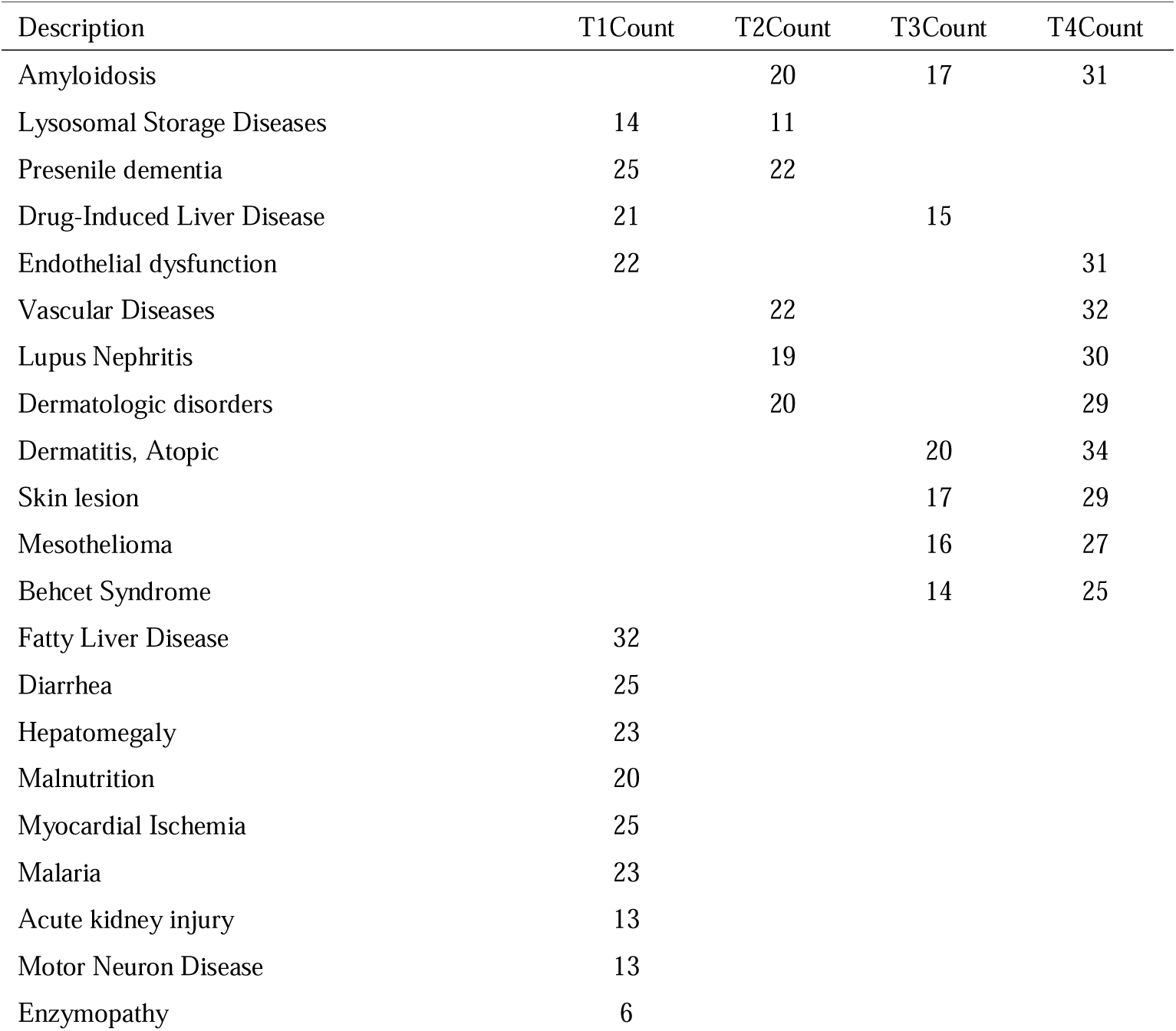
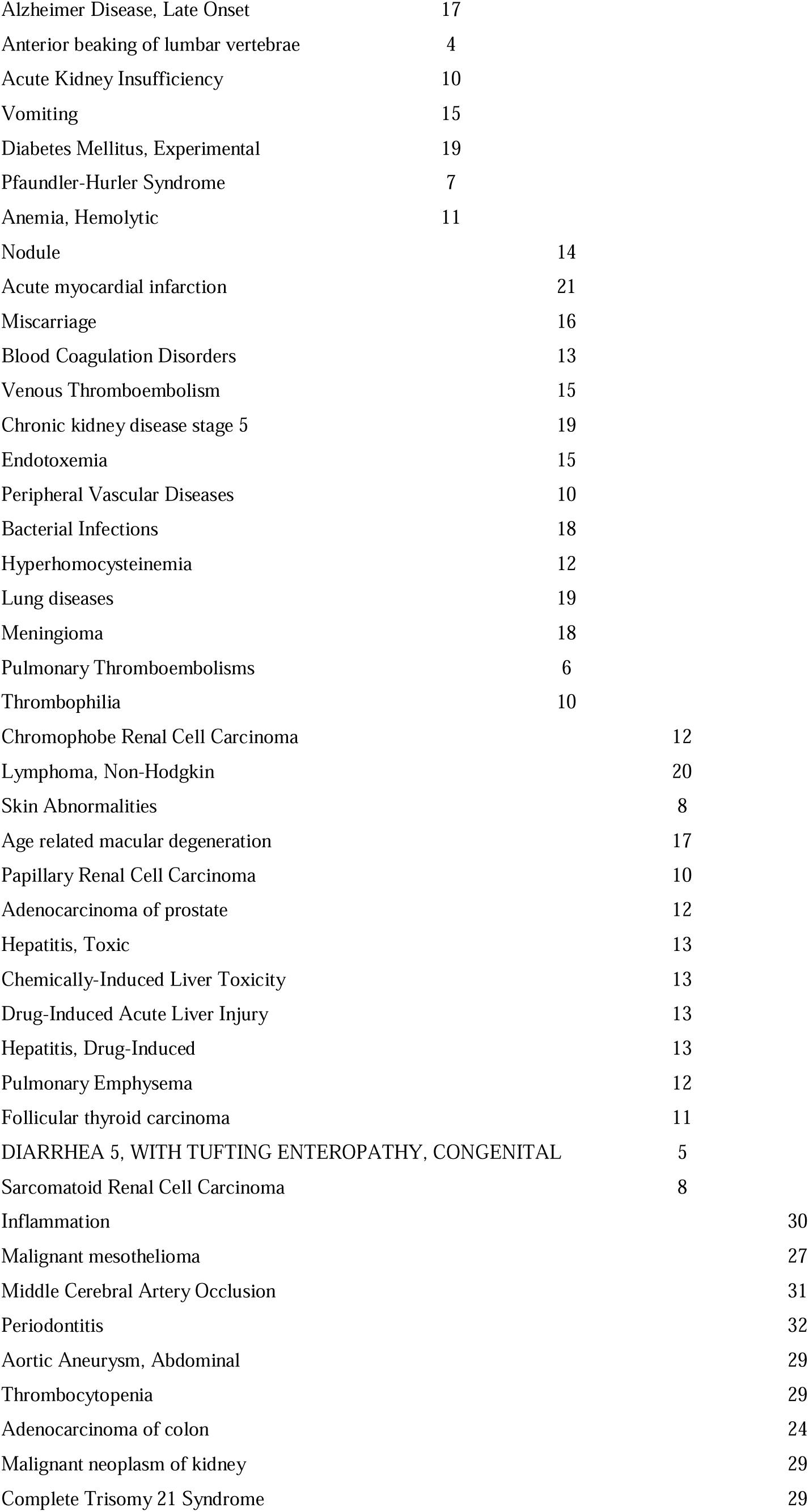
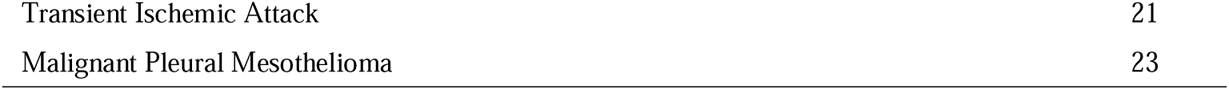
Summary of enrichment analysis in DisGeNET (T1~T4, Homozygote-Heterozygote)

The differential proteins at each time point of T1, T2, and T3 were imported into STRING website for ontology analysis of mammalian phenotype (The Mammalian Phenotype Ontology, Monarch). This function relies on the Monk website. The Monk Initiative is a platform for data integration and analysis, which links phenotype with cross-species genotype, and basic and applied research with semantic-based analysis. Exploring the correlation between phenotypic results, diseases and genetic variation as well as environmental factors, the platform created a biological ontology based on this, which together realized the computational analysis of complex and semantic integration of gene, genotype, variant, disease and phenotypic data, identified animal models of human diseases through phenotypic similarity, provided phenotypically driven computational support for differential diagnosis, and transformation research [16, 17]. Abnormal biological processes or diseases with an FDR<0.01 at each of the 3 time points after phenotypic analysis of the differential protein Monarch are presented in Table 3 below.

**Table 3.**
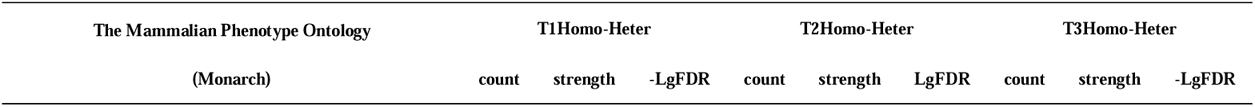
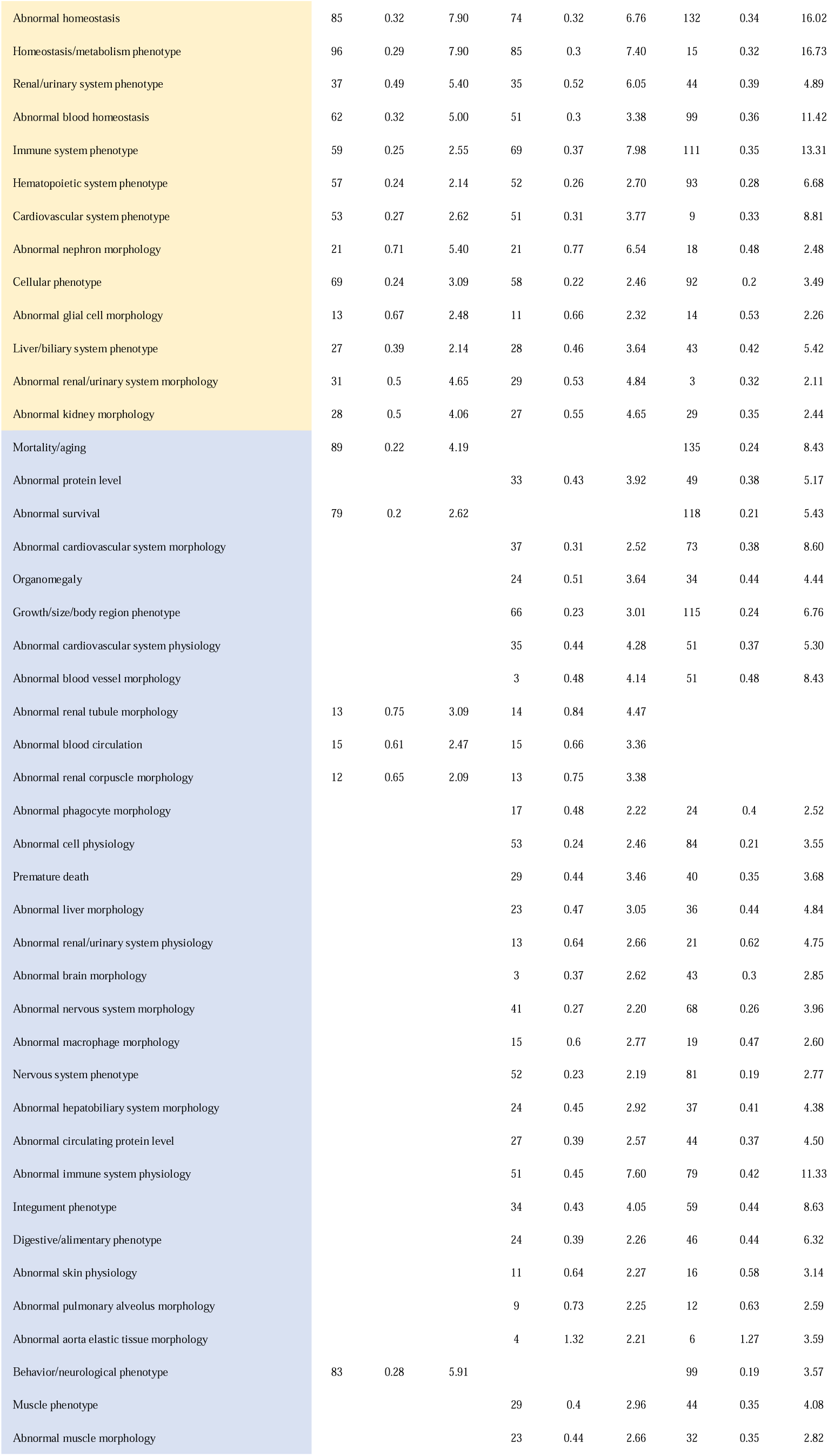
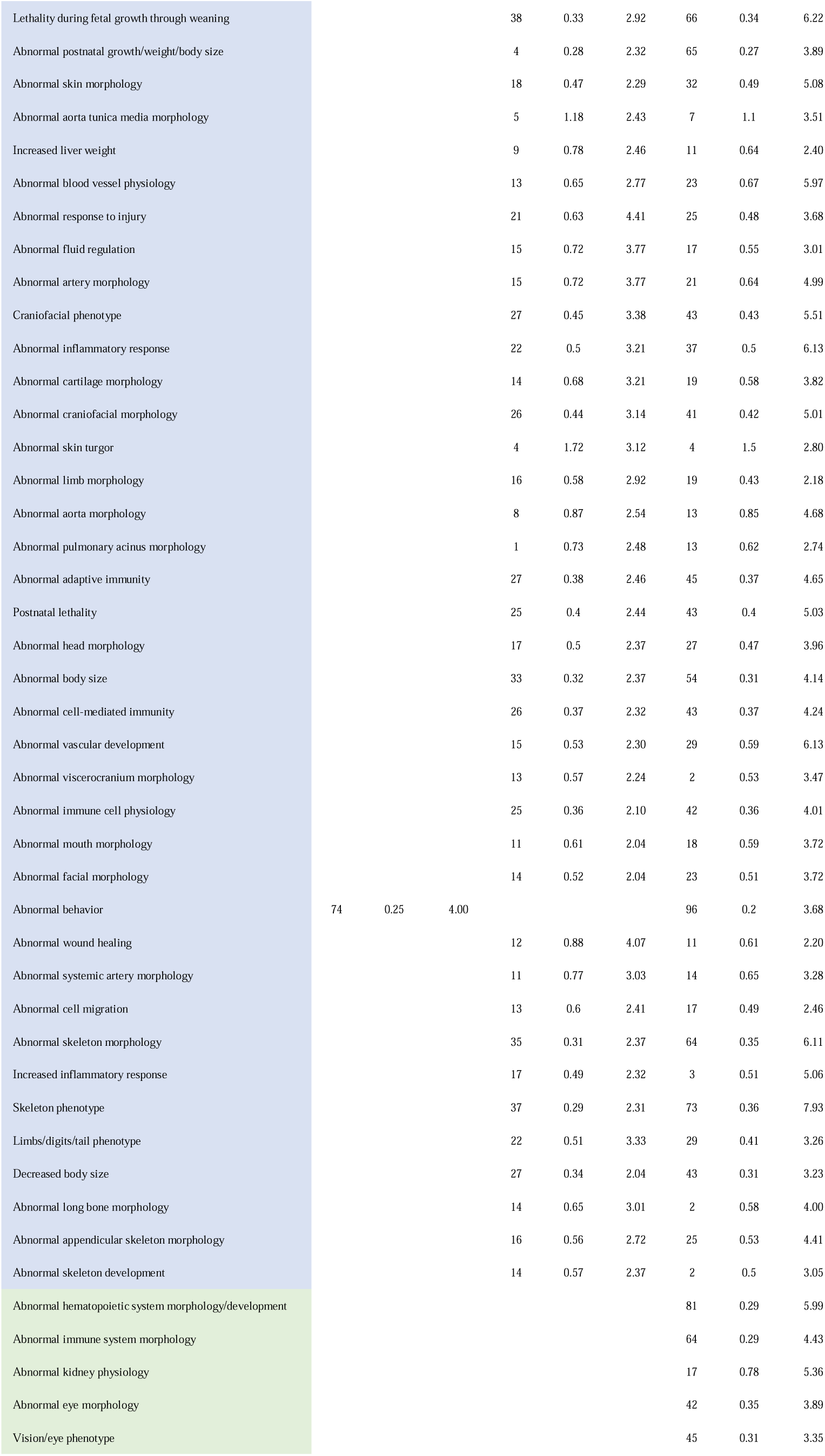
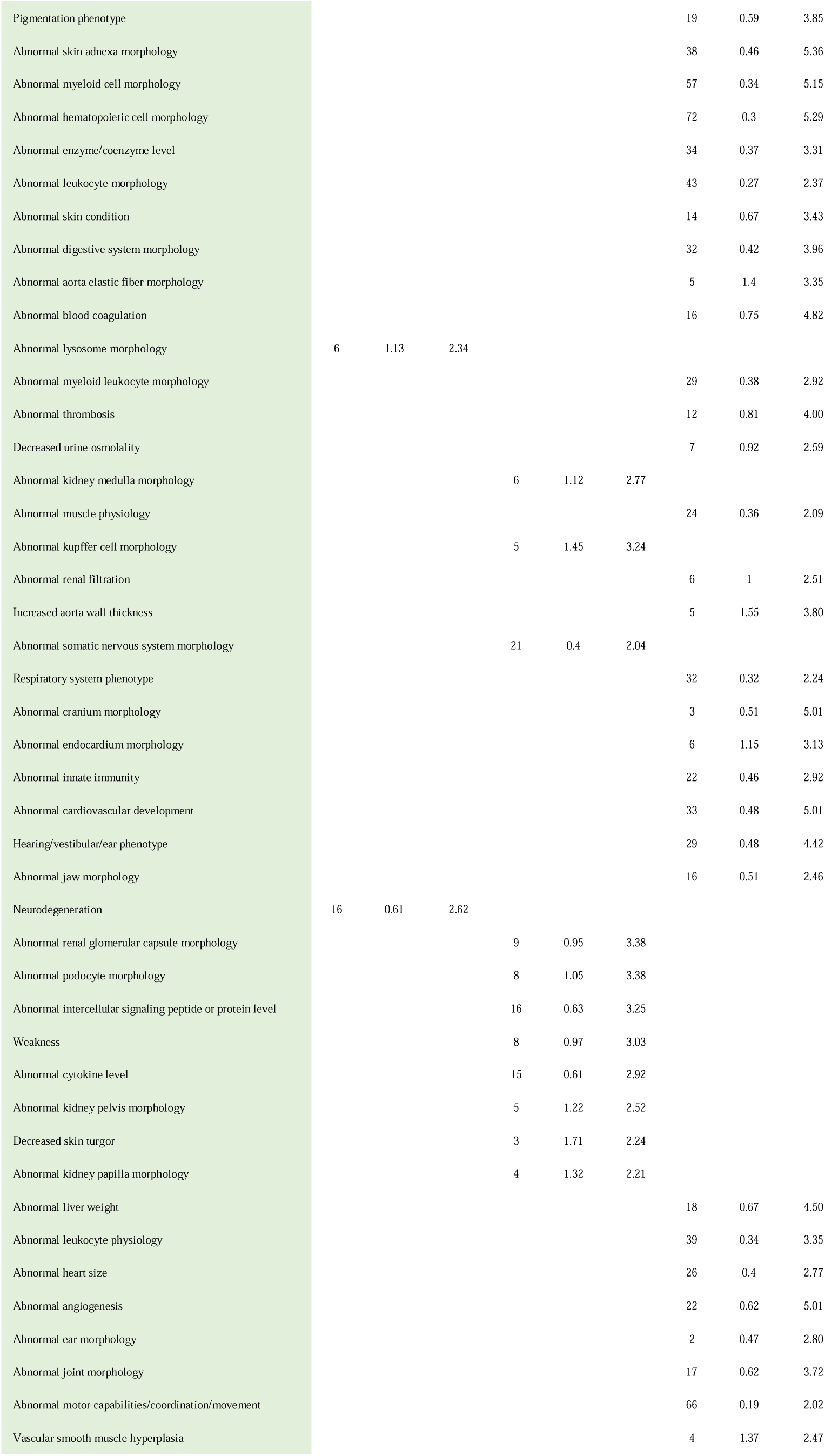
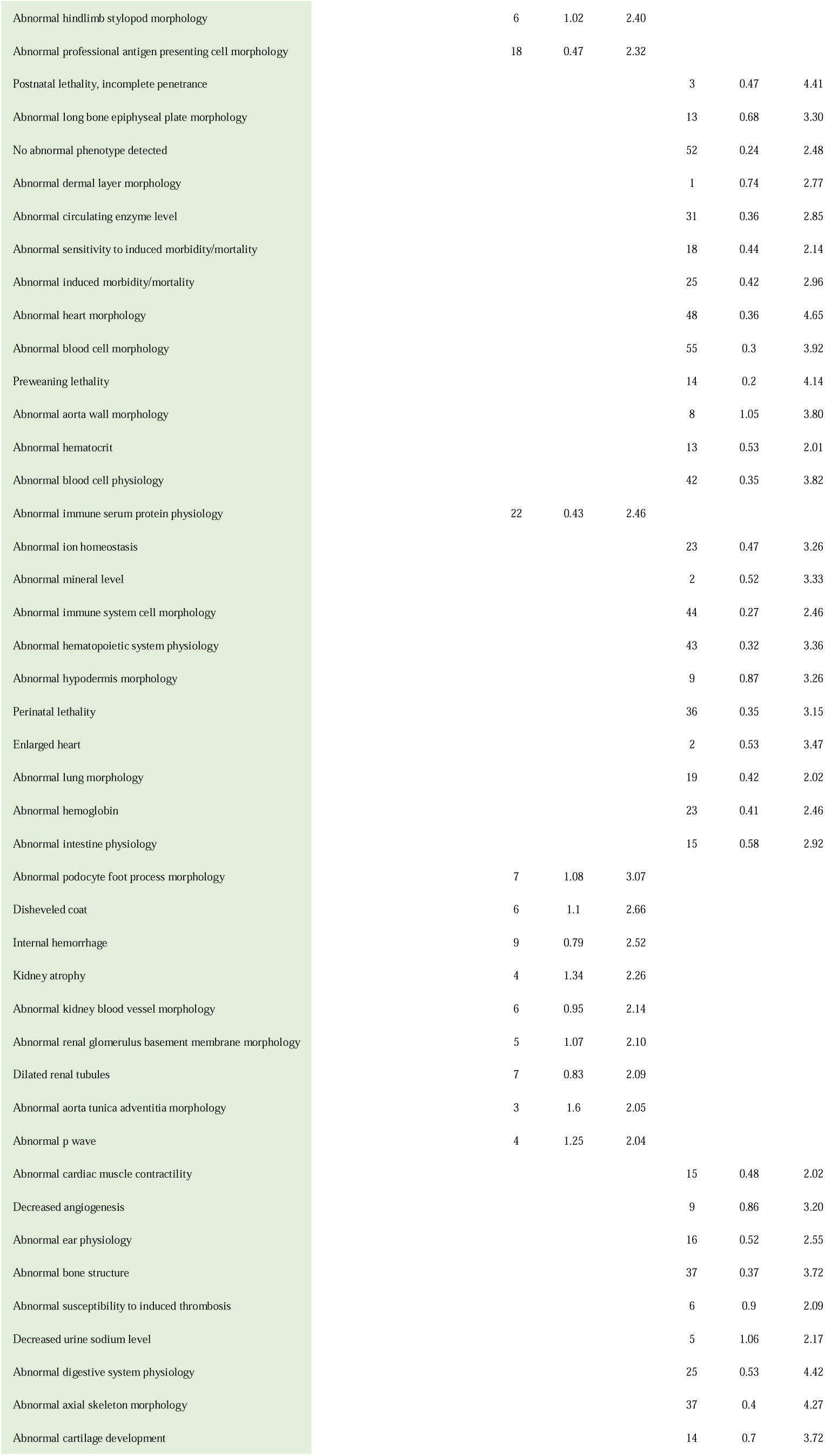
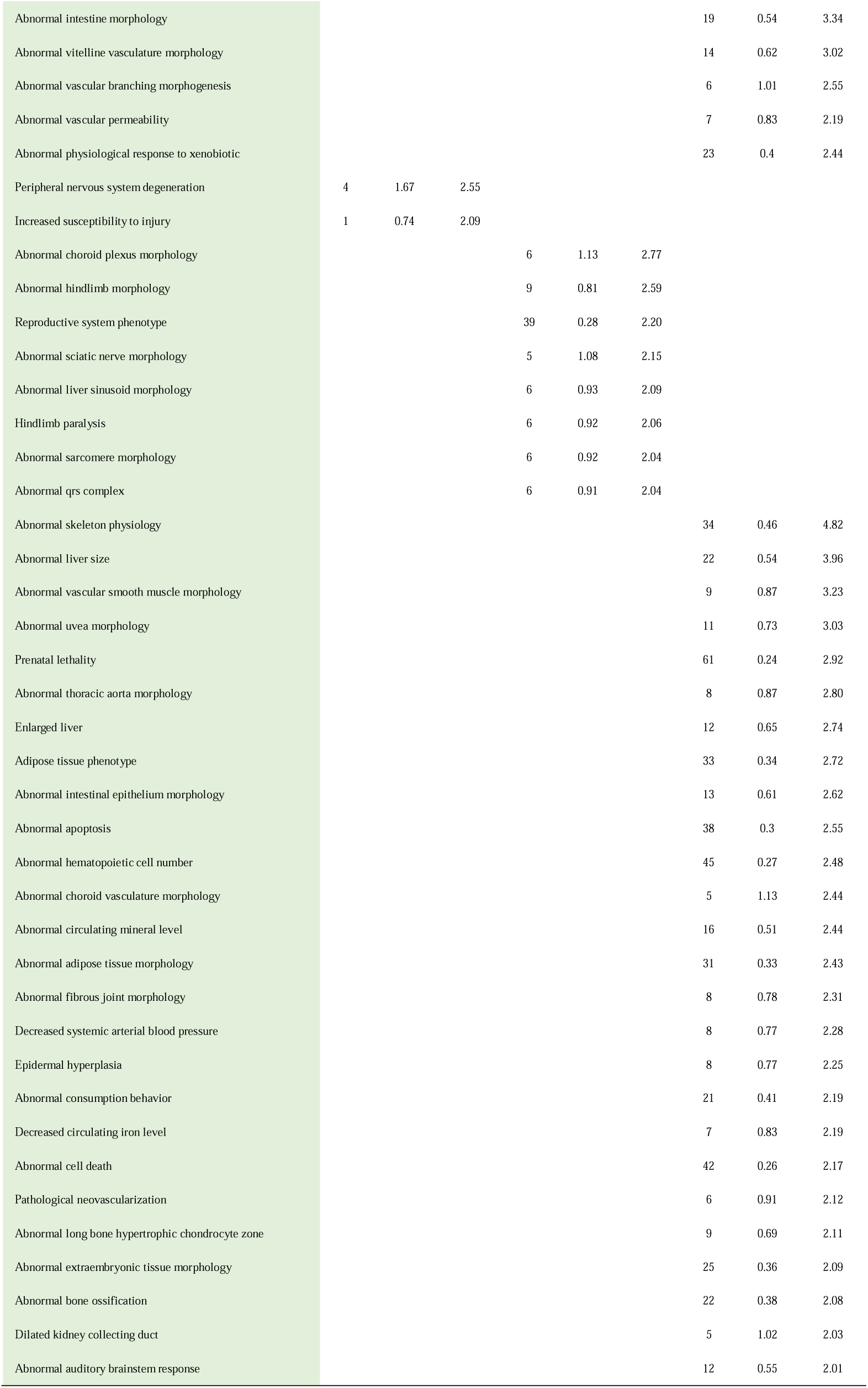
Summary of enrichment analysis in Monarch (T1~T3, Homozygote-Heterozygote)

Common to all time points included abnormal homeostasis, homeostasis/metabolic phenotype, renal/urinary phenotype, abnormal blood homeostasis, immune phenotype, hematopoietic phenotype, cardiovascular phenotype, nephron morphology, cellular phenotype, glial cell morphology, liver/gallbladder phenotype, renal/urinary phenotype, renal morphology. Premature death was found at both T2 and T3 time points, which did not contain identical protein and eight proteins in total. One tumor necrosis factor receptor (Tumor necrosis factor receptor superfamily member 11A) was worth mentioning, and there were 21 and 32 protein differences at T2 and T3 time points, respectively. And the abnormal phenotype involving the two time points further includes, Abnormal protein level, abnormal survival state, cardiovascular system abnormalities, organ hypertrophy, renal tubules and renal corpuscles morphological abnormalities, blood circulation abnormalities, phagocyte morphological abnormalities, liver morphological abnormalities, brain morphological abnormalities, neurological morphological abnormalities, macrophage morphological abnormalities, hepatobiliary morphological abnormalities, immune system abnormalities, skin physiological abnormalities, alveolar morphological abnormalities, aortic intima morphological abnormalities, abnormal inflammatory response, cartilage morphological abnormalities, craniofacial morphological abnormalities, Abnormal skin swelling, limb morphology, head morphology, body shape, oral morphology, facial morphology, behavior, wound healing, cell migration, bone morphology, bone development, etc. There were fewer abnormal (FDR<0.01) pathways unique to T1, including lysosomal morphological abnormalities, neurodegeneration, peripheral nervous system degeneration, and increased body vulnerability. At T2, the number of unique abnormal pathways (FDR<0.01) increased, including renal medulla morphological abnormality, kupffer cell morphological abnormality, somatic nervous system morphological abnormality, glomerular cystic abnormal morphology, podoid process cell morphological abnormality, intercellular signal peptide or protein level abnormality, weakness, cytokine level abnormality, renal pelvic morphological abnormality, renal papilla morphological abnormality, hind limb morphological abnormality, immune serum protein physiological function abnormality, internal hemorrhage, kidney atrophy, renal vascular morphological abnormality, aortic adventitia morphological abnormality, hind limb paralysis, sciatic nerve morphological abnormality, hepatic sinus morphological abnormality, etc. The abnormal (FDR<0.01) pathway unique to T3 was the most. Morphological/developmental abnormalities of hematopoietic system, morphological abnormalities of immune system, renal physiological abnormalities, morphological abnormalities of eyes, skin adnexa, bone marrow cells, hematopoietic cells, enzyme/coenzyme levels, leukocyte morphology, digestive system morphology, aortic elastic fiber morphology, blood coagulation abnormalities, bone marrow leukocyte morphology, thrombosis abnormalities, muscle physiological abnormalities, renal filtration abnormalities, increase in aortic wall thickness, Cranial morphological abnormalities, endocardial morphological abnormalities, innate immune abnormalities, cardiovascular developmental abnormalities, maxillofacial morphological abnormalities, liver weight abnormalities, leukocyte physiological abnormalities, heart size abnormalities, angiogenesis abnormalities, ear morphological abnormalities, joint morphological abnormalities, abnormal motor ability/coordination/movement, long epiphyseal plate morphological abnormalities, dermal layer morphological abnormalities, abnormal levels of circulating enzymes, susceptibility to induction of morbidity and mortality abnormalities, heart morphological abnormalities, blood cell morphological abnormalities, Abnormal aortic wall morphology, abnormal blood cell physiological function, abnormal ion homeostasis, abnormal mineral level, abnormal immune system cell morphology, abnormal hematopoietic system physiological function, abnormal subcutaneous tissue morphology, abnormal lung morphology, abnormal hemoglobin, abnormal intestinal physiological function, abnormal myocardial contractility, decreased angiogenesis, abnormal ear physiological function, abnormal bone structure, abnormal susceptibility to induce thrombosis, abnormal digestive system physiological function, abnormal axial bone morphology, Cartilage dysplasia, intestinal morphological abnormalities, yolk blood vessel morphological abnormalities, vascular branch morphology abnormalities, vascular permeability abnormalities, bone physiological function abnormalities, liver size abnormalities, vascular smooth muscle morphological abnormalities, thoracic aorta morphological abnormalities, liver enlargement, intestinal epithelial morphological abnormalities, abnormal apoptosis, abnormal number of hematopoietic cells, choroidal blood vessel morphological abnormalities, abnormal levels of circulating minerals, adipose tissue morphological abnormalities, fibrous joint morphological abnormalities, systemic arterial blood pressure drop, Epidermal hyperplasia, decreased circulating iron levels, abnormal cell death, pathological neovascularization, morphological abnormalities of extra-embryonic tissue, abnormal ossification of bone, dilated renal collecting duct, abnormal auditory brainstem response.

### Comparative analysis between homozygous/heterozygous and wild type at the same time point

The differences between homozygote or heterozygote and wild-type at each time point were analyzed, and the differential proteins were screened according to the standard that the P value was < 0.05 and the fold-change value was < 0.5 or > 2. The differential proteins at T1, T2, T3 and T4 at each time point were then imported to the Metascape website for enrichment function analysis. A more significant pathway for homozygous and wild-type differential protein enrichment to the top 20 is shown in Table 4 below, set up as described in the previous section, using methods such as Kegg Pathway, Go Biological Procedures, Reach OME Gene Sets, Canonical Pathways, Corum, WikiPathways et al. Ontology and PANTHER Pathway, all the entered differential proteins were used as the background for enrichment analysis, limiting the p-value for each pathway process to < 0.01, the minimum count to 3, and the enrichment factor to > 1.5.

**Table 4.**
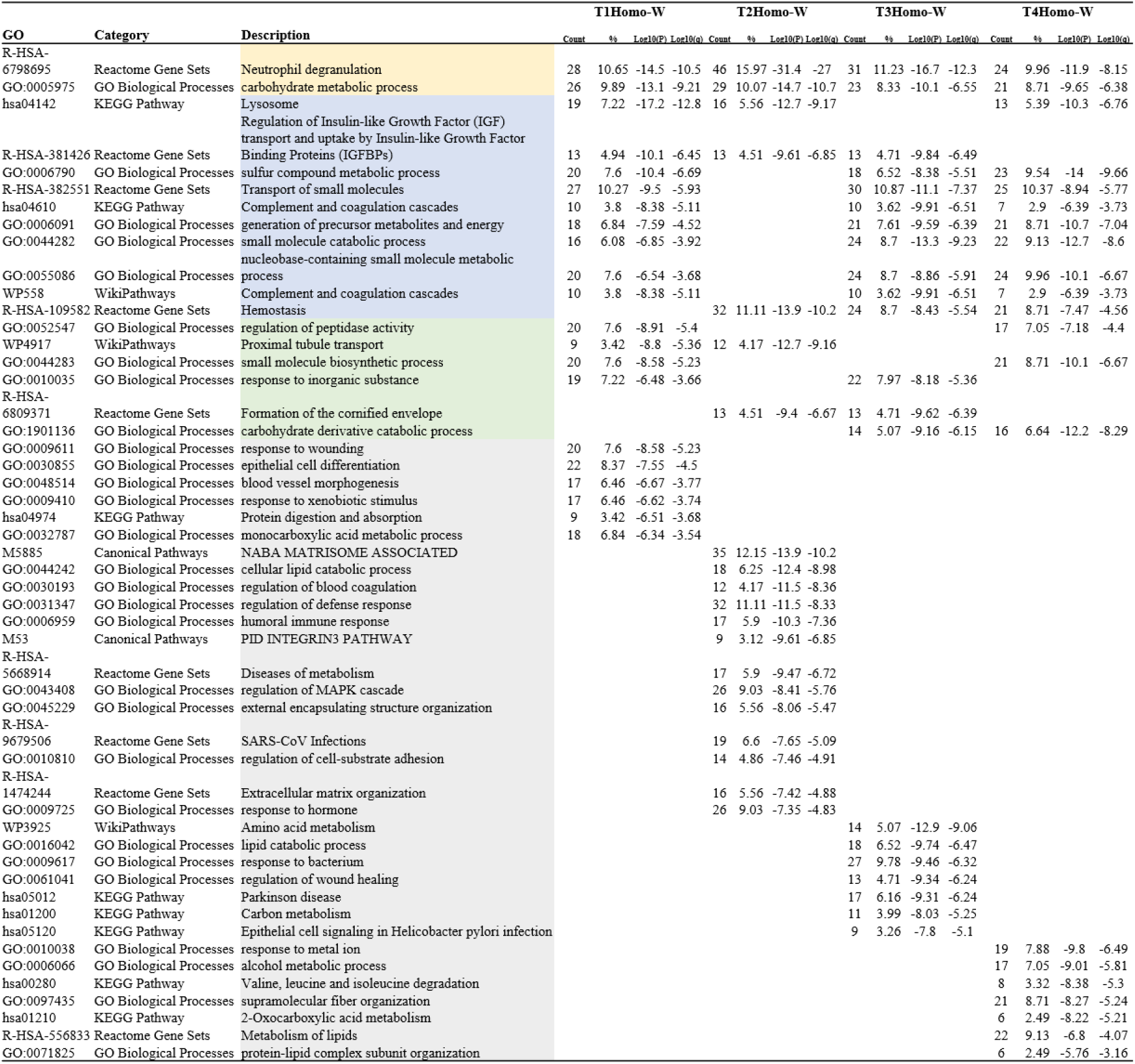
Top 20 clusters with their representative enriched biological processes terms (one per cluster, T1~T4 Homozygote-WT)

At the four time points, homozygous and wild-type differential protein enrichment shared two pathways, namely, neutrophil degranulation and carbohydrate metabolism. The three time points involved were lysosomes, regulation of insulin-like growth factor binding protein on insulin-like growth factor (IGF) transport and uptake, metabolic processes of sulfur-containing compounds, transport of small molecules, complement and coagulation cascade, production of precursor metabolites and energy, small molecule catabolism, small molecule metabolism with nucleobases, complement and coagulation cascade, and hemostasis. Regulation of peptidase activity, proximal tubular transport, small molecule biosynthesis processes, response to inorganic matter, formation of keratinized envelopes, and catabolism of carbohydrate derivatives were involved at two time points. The unique pathways for each time point were as follows: T1: response to trauma, epithelial cell differentiation, vascular morphogenesis, response to xenobiotic stimulation, protein digestion and absorption, and monocarboxylic acid metabolism; T2: NABA stroma complex, cellular liPID catabolism process, regulation of blood coagulation, regulation of defense response, humoral immune response, pid integrin 3 pathway, metabolic disease, regulation of MAPK cascade, external packaging structure tissue, SARS coronavirus infection, regulation of cell-matrix adhesion, extracellular matrix tissue, response to hormones; T3: amino acid metabolism, lipid catabolism processes, reaction to bacteria, regulation of wound healing, parkinson’s disease, carbon metabolism, epithelial signaling in helicobacter pylori infection; T4: Reaction to metal ions, alcohol metabolism, valine, leucine and isoleucine degradation, supramolecular fiber polymerization, 2-oxocarboxylic acid metabolism, lipid metabolism, protein-lipid complex subunit composition. Although the specific pathways at each time point were different, they all included metabolic-related pathways, so it could be seen that p53 gene did have a greater impact on metabolism.

Differential proteins obtained by comparing homozygosity and wildtype at each time point were analyzed for gene and disease associations by DisGeNET. DisGeNET is a discovery platform that contains the largest publicly available collection of genes and their variants related to human diseases [18–20]. DisGeNET consolidated data from a database of professional organizations, the GWAS catalog, animal models, and the scientific literature. Distributed network data is uniformly annotated with controlled vocabularies and community-driven ontologies. In addition, several primary indicators were provided to help prioritize the genotypic-phenotypic relationship. The current version of DisGeNET (v7.0) contains 1134942 gene-disease associations (GDAs) between 21671 genes and 30170 diseases, disorders, traits and clinical or abnormal human phenotypes, and 369554 mutation-disease associations (VDAs) between 194515 mutations and 14155 diseases, traits and phenotypes. Table 5 below shows a series of top-ranked related diseases, with no disease phenotype in common at the four time points, and the diseases in common at the three time points as follows: lung disease, chronic kidney disease stage 5, diabetes, and lupus nephritis; Diseases common to both time points were as follows: emphysema, malformed osteoarthropathy, fatty liver, pneumonia, atherosclerotic lesions, acute myocardial infarction, carotid atherosclerosis, vascular diseases, venous thromboembolism, non-filtering intraductal carcinoma, meningioma; Diseases unique to each time point were as follows: T1: congenital pulmonary hypertension, hypercholesterolemia, behcet’ s syndrome, ankylosing spondylitis, follicular adenoma, down’ s syndrome, chromophobe cell carcinoma of the kidney, endothelial dysfunction, trisomy 21 syndrome, meningitis, lysosomal storage disease, T2: respiratory distress syndrome, follicular thyroid canc, fibrosis, cancer spread, thrombophilia, hepatomegaly, oral malignancy, papillary renal cell carcinoma, canc transitional cells, benign prostatic hyperplasia, thrombocytopenia, T3: hyperhomocysteinemia, encephalopathy, renal cancer, congenital diaphragmatic hernia, sarcomatosis, left heart failure, lymphoma, Non-Hodgkin, cerebral cavernous hemangioma, blindness, intercellular degeneration, secondary malignant tumor of bone, fatigue, scleroderma, T4: metabolic/homeostatic abnormalities, abdominal aortic aneurysm, cardiac arrest, hyperuricemia, amyloidosis, pre-senile dementia, vomiting, middle cerebral artery occlusion, mental decline, primary thrombocytosis, hyperlipidemia. Mental decline or mental decline is one of the characteristics of neurodegenerative diseases [21]. In general, the accumulated diseases include more lung-related diseases, many cardiovascular-related diseases, tumors or cancers, and of course, more neurodegenerative-related diseases.

**Table 5.**
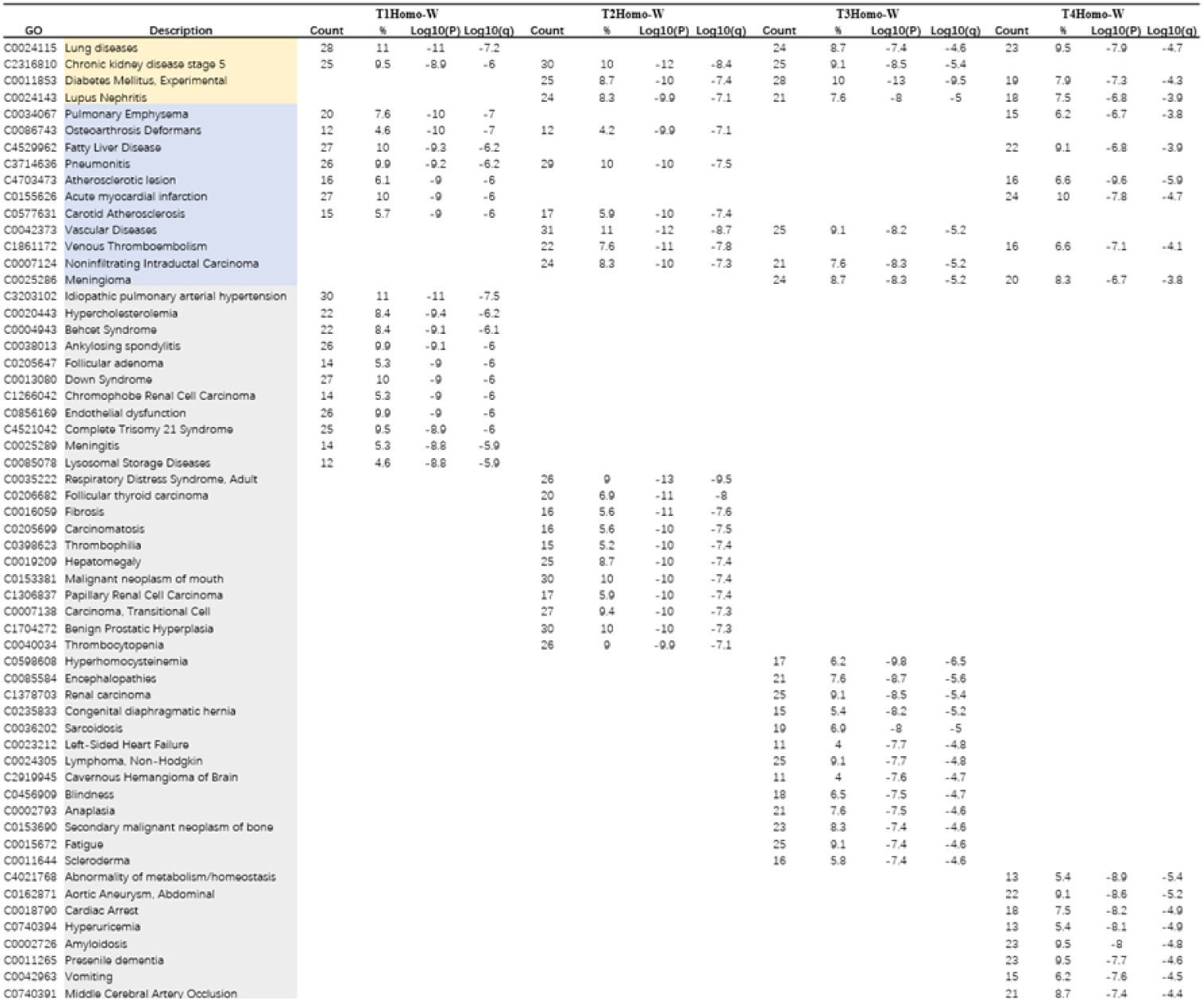
Top 20 clusters with their representative enriched related disease terms (one per cluster, T1~T4 Homozygote-WT)

The differential proteins obtained by comparing the heterozygotes with the wild-type at each time point were analyzed for enrichment function by Metascape website, and the main biological processes in the top 20 of each of the four time points are shown in Table 6 below. Biological processes common to the four time points included: small molecule catabolism, neutrophil degranulation, response to metal ions, sulfur-containing compound metabolism, and response to exogenous compound stimulation. Biological processes common to the three time points included: small molecule biosynthesis, carbohydrate metabolism, precursor metabolite and energy production, nucleobase-containing small molecule metabolism, lysosomes, response to toxic substances, small molecule transport, and hemostasis. Biological processes common to the two time points included monocarboxylic acid metabolism, alcohol metabolism, intercellular adhesion, extracellular matrix tissue, regulation of peptidase activity, lipid catabolism, amino acid metabolism, and metabolism of vitamins and cofactors. The specific biological processes at T1 were: regulation of insulin-like growth factor (IGF) transport and uptake by IGF-binding protein, glycolysis/gluconeogenesis, degradation of valine, leucine and isoleucine, and supramolecular fibrous tissue; Biological processes specific to the T2 time point were: sulfur amino acid metabolism, VEGFA-VEGFR2 signaling pathway (VEGF regulates tumor angiogenesis), proximal tubular transport, PID HIF1 TFPATHWAY, transferrin endocytosis and recirculation, purinergic compound metabolism, actin cytoskeleton regulation, epithelial cell differentiation, response to bacteria, complement and coagulation cascades, and regulation of vesicle-mediated transport. The specific biological process at T3 was: response to cadmium. Biological processes specific to the T4 time point were negative regulation of hydrolase activity, keratinocyte differentiation, reaction to alcohol, and Parkinson’s disease. Comparative analysis of heterozygotes and wild-type revealed that the biological pathways enriched were still dominated by metabolism. It is worth noting that the last time point was enriched for Parkinson’s disease, an senile degenerative neurological disease, while homozygotes were enriched for Parkinson’s disease at time point T3, which may be one of the reasons for the premature death of p53 knockout mice as a whole and the death of homozygotes before heterozygotes.

**Table 6.**
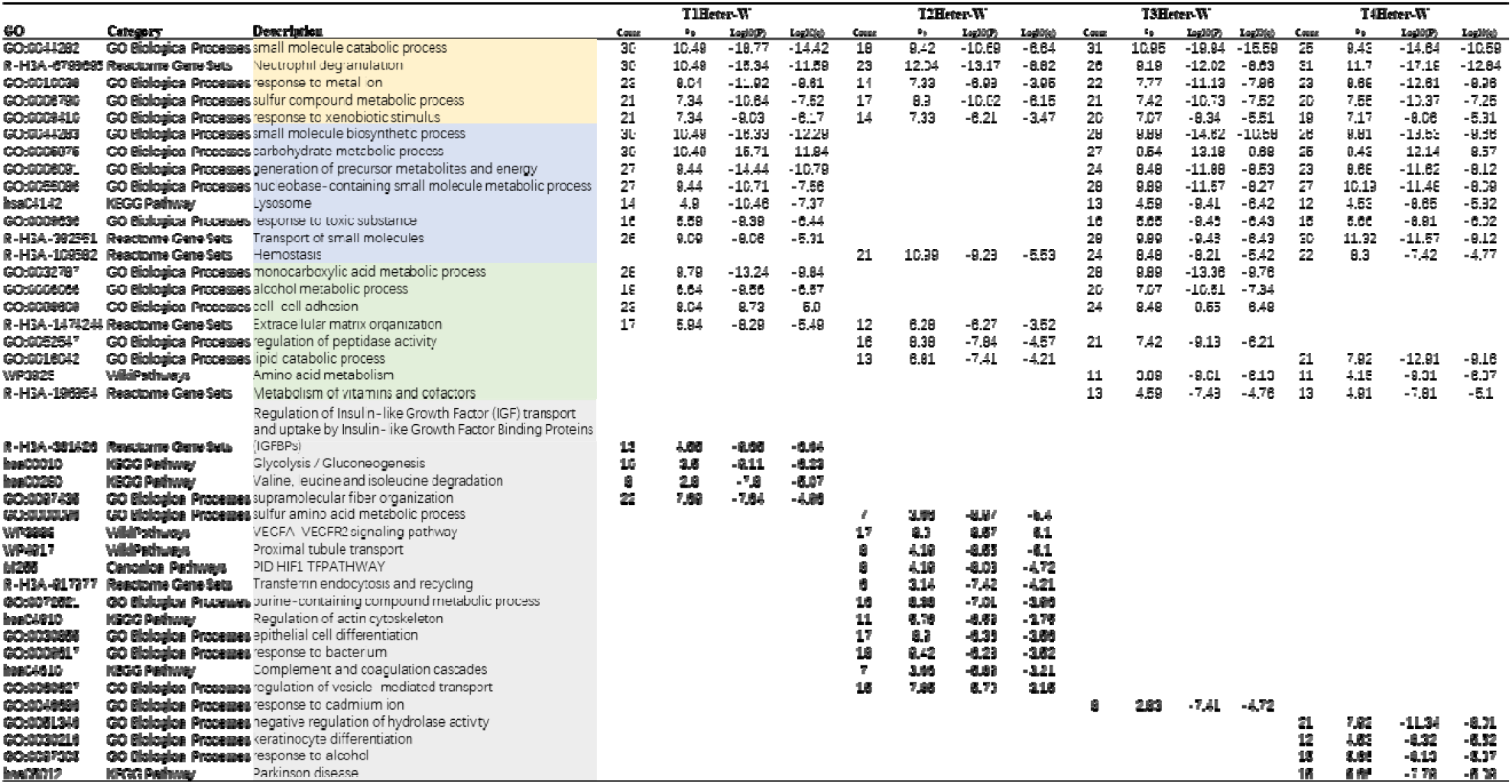
Top 20 clusters with their representative enriched biological processes terms (one per cluster, T1~T4 Heterozygote-WT)

The differential proteins obtained by comparing the heterozygotes with the wild-type at each time point were analyzed for gene and disease association by DisGeNET, as shown in Table 7 below. The common biological pathways at the four time points were found to be fatty liver and drug-induced liver disease. It can be seen that liver disease is very important in the heterozygous association, which differs from homozygosity in that lung disease comes first in homozygosity. The disease pathways common to the three time points include: lung disease, retinal disease, atherosclerotic lesion, diabetes, senile dementia, cancer spread, pancreatitis, and adult respiratory distress syndrome, such as lung disease, atherosclerotic lesion, diabetes, senile dementia, and cancer spread. These diseases can be enriched in the first time point, suggesting that urine may reflect the early stage of the disease. Biological pathways of disease involving two time points included: carotid atherosclerosis, hyperuricemia, acute coronary syndrome, malignant pleural mesothelioma, meningioma, emphysema, acute myocardial infarction, and undifferentiated carcinoma. For example, malignant pleural mesothelioma is enriched at the first time point, meningioma begins at the third time point, and undifferentiated cancer begins at the second time point. Undifferentiated means that the biological behavior of cancer cells is completely different from the normal tissues from which they originate. They are in a very primitive state, and it is sometimes even difficult to identify the tissue source of undifferentiated cancer cells. Therefore, undifferentiated cancer refers to cancer cells that are very primitive and have a high degree of malignancy. As far as we know, some early undifferentiated cancers can still be cured after active surgical treatment, radical radiotherapy and chemotherapy, and the cure rate can even reach more than 50%. For some patients who belong to undifferentiated cancer and stage it later, the therapeutic effect is relatively poor and it is difficult to obtain a good therapeutic effect, let alone the statement that there is no cure. However, the pathways leading to the undifferentiated cancer can be enriched in the second time point, i.e., 42 days after lactation. Perhaps by utilizing the sensitivity of urine, the malignant cancer can be found earlier through the early diagnosis of urine, thus improving the survival rate and curative rate of people. Each time point also had unique disease pathways enriched therein. Time points T1 included: chemical and drug-induced liver injury, hepatitis, chemical liver toxicity, drug-induced acute liver injury, drug-induced hepatitis, vascular disease, primary myelofibrosis, and lupus nephritis. It could be seen that most of the diseases in the liver at this early time point were related to diseases. T2 time points include: mental decline, venous thromboembolism, pancreatic tumor, mesothelioma. Mental decline is one of the signs of neurodegenerative diseases, and tumor and thrombus have appeared in the second time point. Time points T3 included chronic pancreatitis, alcoholic pancreatitis, abdominal aortic aneurysm, pneumonia, anemia, hypertriglyceridemia, memory disorders, hyperlipidemia, and memory disorders that may have contributed to the abnormal aging of the mouse body and eventually to premature death. T4 time points include: undifferentiated cancer deformability, osteoarthropathy, breast tumor, breast cancer, spindle cell carcinoma, follicular adenoma, colonic adenocarcinoma, obstructive sleep apnea, irritable bowel syndrome, primary thrombocytosis, papillary renal cell carcinoma, and familial idiopathic cardiomyopathy. It can be seen that many cancers appeared at the last time point, which is much more obvious than that at the previous time point. This also indicates that the possibility of tumor cancer induction after p53 knockout is higher, which is also one of the reasons for the early death of the mice.

**Table 7.**
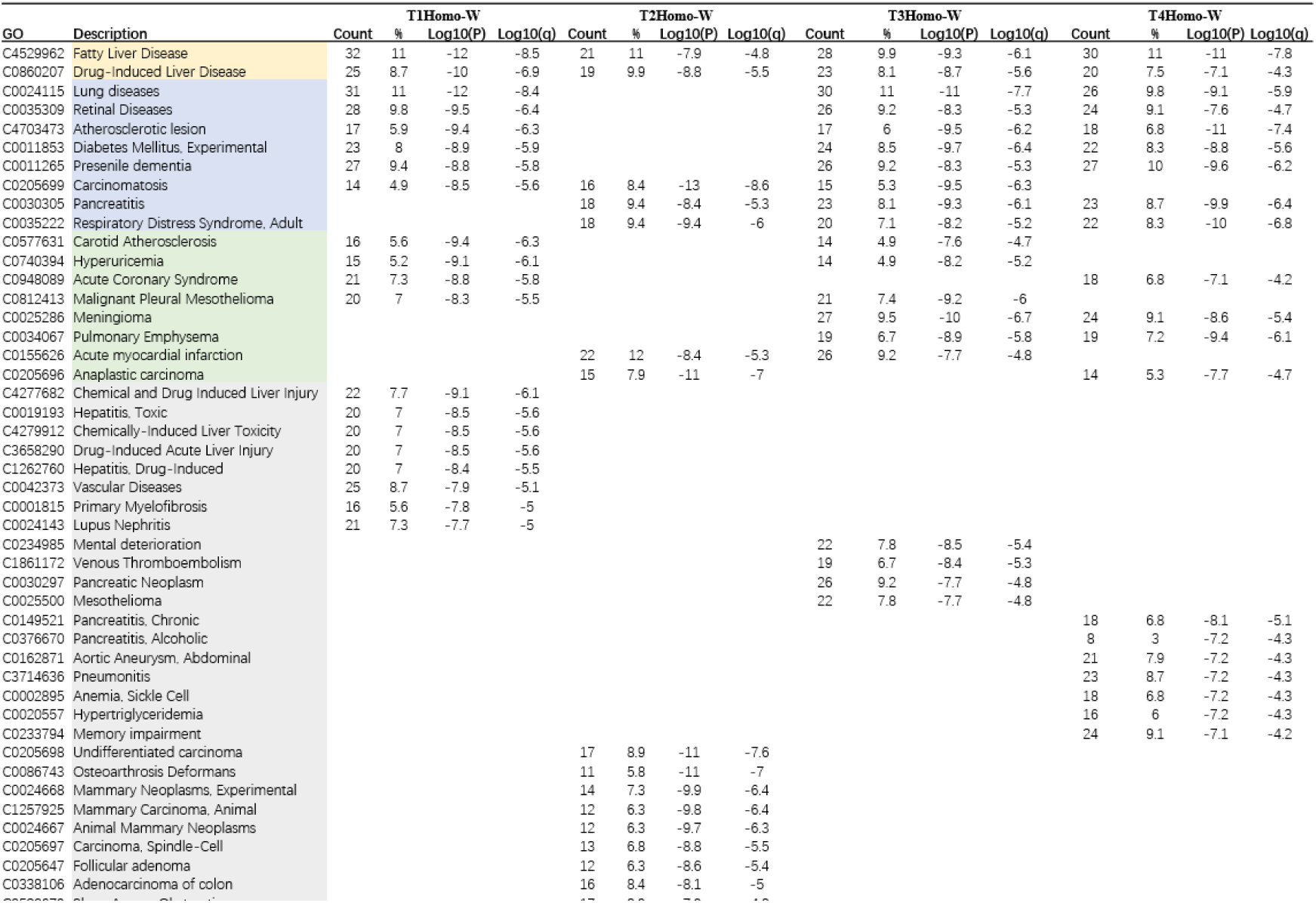
Top 20 clusters with their representative enriched related disease terms (one per cluster, T1~T4 Heterozygote-WT)

We compared the differential proteins obtained by homozygosity and wild-type as well as the enriched pathways with those obtained by disease and heterozygosity, as shown in Fig. 12 below. The differential proteins at each time point were more or less repetitive, but none of the coincident protein species exceeded the sum of the respective unique differential proteins. In the first time point section, the first Venn diagram showed the crossing of differential proteins between the two, with a total of 180 differential proteins, and homozygous and heterozygous had 116 and 137 unique differential proteins respectively. The second Venn diagram showed the crossing of pathways enriched to the top 20. In T1, biological pathways were mainly metabolized, and there were five metabolism-related processes out of 12 common pathways. Including nucleobase-containing micromolecule metabolism process, neutrophil degranulation, micromolecule biosynthesis process, exogenous hormone stimulation reaction, lysosome, micromolecule transportation, precursor metabolite and energy generation, monocarboxylic acid metabolism process, insulin-like growth factor binding protein regulation on IGF transport and uptake, sulfur-containing compound metabolism process, carbohydrate metabolism process and micromolecule catabolism process; In addition, the third Venn diagram shows the intersection of the two diseases, with 4 disease phenotypes including carotid atherosclerosis, pulmonary disease, fatty liver and atherosclerotic lesion. In the second time point, there were 96 proteins in total; 229 proteins were unique to homozygote and 115 proteins were unique to heterozygote; and there were four overlapping pathways, namely, extracellular matrix tissue, hemostasis, proximal tubular transport, and neutrophil degranulation. There were four overlapping diseases, namely, papillary renal cell carcinoma, malformed osteoarthropathy, carcinoma spread, and respiratory distress syndrome. In the third time point, they shared 162 proteins, with 147 and 153 unique proteins for homozygote and heterozygote respectively, and 9 overlapping pathways, including small molecule metabolism process containing nucleobases, degranulation of neutrophils, small molecule transportation, production of precursor metabolites and energy, amino acid metabolism, sulfur-containing compound metabolism process, carbohydrate metabolism process, hemostasis, and small molecule catabolism process. All three diseases were related, i.e., lung disease, meningioma, and diabetes. In the fourth time point, they shared 181 proteins, 96 unique proteins for homozygote and 116 unique proteins for heterozygote, and 11 overlapping pathways. Including nucleobase-containing micromolecule metabolism process, neutrophil degranulation, micromolecule biosynthesis process, lysosome, micromolecule transportation, precursor metabolite and energy generation, sulfur-containing compound metabolism process, carbohydrate metabolism process, hemostasis, response to metal ions and micromolecule catabolism process. To sum up, biological pathways are mainly concentrated in metabolism and neutrophil degranulation, while diseases are mainly concentrated in cardiovascular and cerebrovascular diseases, cancer and tumor, and lung and liver diseases. The mating situations of the differential proteins obtained by comparing the homozygote/heterozygote with the wild type at the four time points are shown in Fig. 13 below. The picture on the left shows the intersection of the differential proteins of the homozygote and the wild type, and there were 66 differential proteins at the four time points, with the coincidence situation significantly more than that of the homozygote and the heterozygote. Each time point also had its own unique differential protein. The picture on the right is the intersection of differential proteins of heterozygote and wild type. There were 120 differential proteins in total at the four time points, and the coincidence was more obvious, indicating that the change of heterozygote with time was slightly slower than that of homozygote, and also indicating that heterozygote was less affected than homozygote after p53 gene was knocked out.

**Fig. 12.**
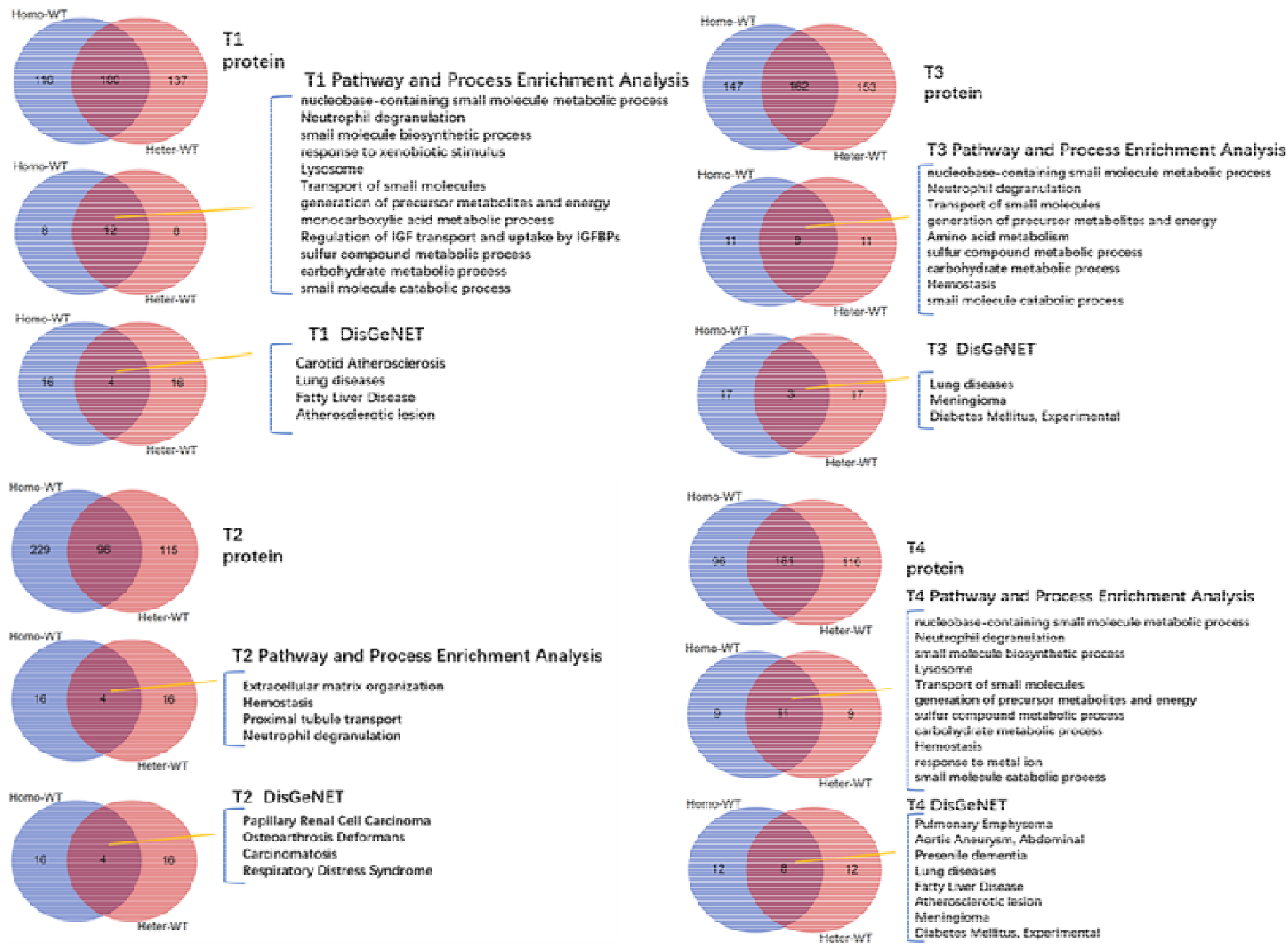
The overlap of differential proteins, biological pathways and diseases of homozygotes and heterozygotes in T1~T4.

**Fig. 13.**
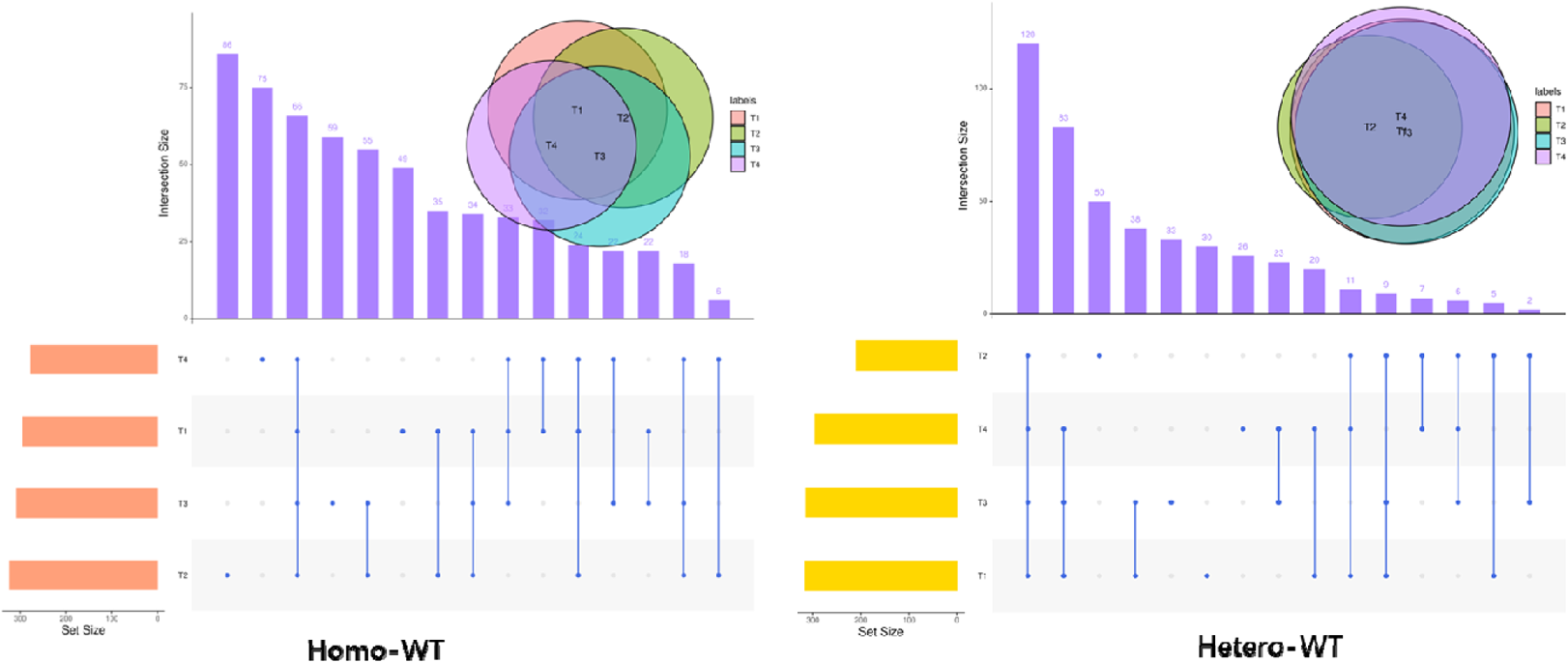
The interaction between homozygote/heterozygote and wild-type differential proteins at four time points

### Self-control analysis of homozygote/heterozygote

Previously, we analyzed homozygote and heterozygote, homozygote and wild-type as well as heterozygote and wild-type respectively at the same time point and found related pathways or gene phenotypes such as cancer, tumor, cardiovascular and cerebrovascular diseases, and neurodegenerative diseases. Next, we performed comparative analysis of homozygote/heterozygote at the front and rear time points. There are six forms of comparison in arrangement and combination at the four time points. Also, the differential proteins were screened according to the criteria with p value < 0.05 and fold-change value < 0.5 or > 2, and then the differential proteins obtained from the comparison at every two time points were imported to the Metascape website for enrichment function analysis. In comparative analysis of homozygosity at two time points, we found that the pathways enriched to the top involved many metabolic processes, as shown in Table 8 below, including production of prerequisite metabolites and energy, small-molecule catabolism, sulfur compound metabolism, aldehyde metabolism in cells, monocarboxylic acid metabolism, carbohydrate metabolism, prostaglandin metabolism, carbon metabolism, drug metabolism, peptide metabolism, carbohydrate derivative catabolism, small-molecule metabolism containing nucleobases, organic hydroxyl compound metabolism, vitamins and cofactors metabolism, and cellular ketone metabolism. In addition, there are also stress, regulatory mediation and other biological processes. Meanwhile, the differential proteins under each comparison were analyzed for gene and disease association by DisGeNET, as shown in Table 9 below. We can find that lupus nephritis exists no matter how the comparison is made at the time point. And most of the time points such as respiratory distress syndrome, adult meningioma, vascular disease, pulmonary disease, pneumonia, diabetes, lymphoma, amyloidosis, skin damage, mesothelioma, fatty liver. However, most of these diseases occurred in the first two time points, indicating that the body condition of p53-knockout mice deteriorated soon after birth, with the appearance of cardiovascular and cerebrovascular diseases, amyloidosis, and tumor cancer. However, the phenotypes of undifferentiated cancer, cancer spread, skin tumor, colon adenocarcinoma, breast tumor, and nephrolithiasis already appeared at the second time point, as well as anemia, Down’s syndrome, middle cerebral artery occlusion and lip cancer, oral cancer, hyperuricemia, and drug-induced liver disease. Diseases unique to T2 compared with T1 are enriched to malignant glioma, skin disease, transitional cell carcinoma, acute pancreatitis, non-ICD-O subtype adult meningioma, and breast cancer. Presenile dementia, myopathy, atherosclerotic lesions, myocardial ischemia, muscle weakness, diabetic complications, motor neurons, delayed Alzheimer’s disease, chronic liver disease, liver injury, and chiropractic-carpal-osseous combination syndrome were enriched at the last time point compared to the first time point (T4T1). It can be seen that senile neurodegenerative diseases have appeared on the 70th day, together with muscle weakness and diabetic complications, which may presage abnormal aging and aggravation of the body until early death.

**Table 8.**
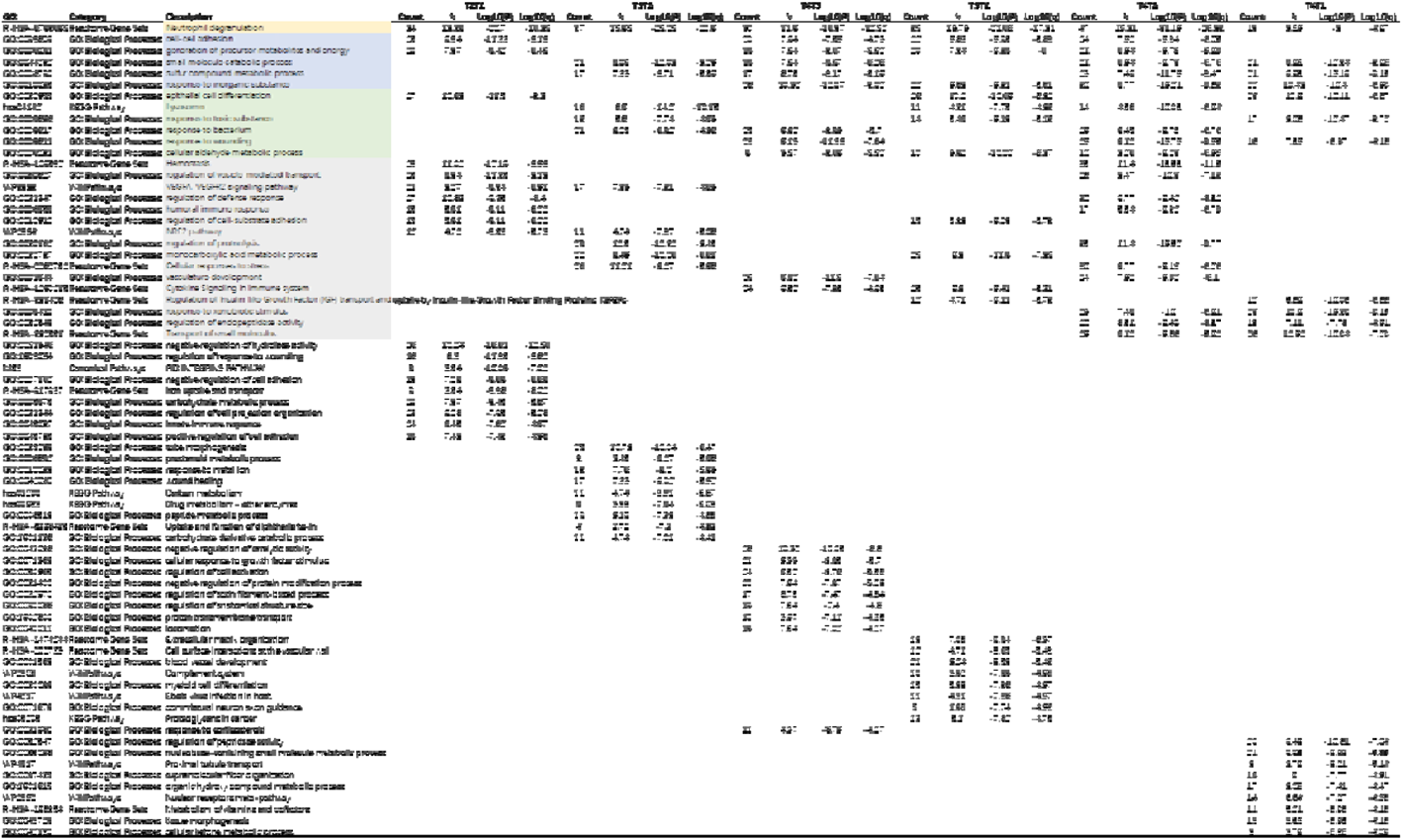
Top 20 clusters with their representative enriched biological processes terms before and after homozygous time points

**Table 9.**
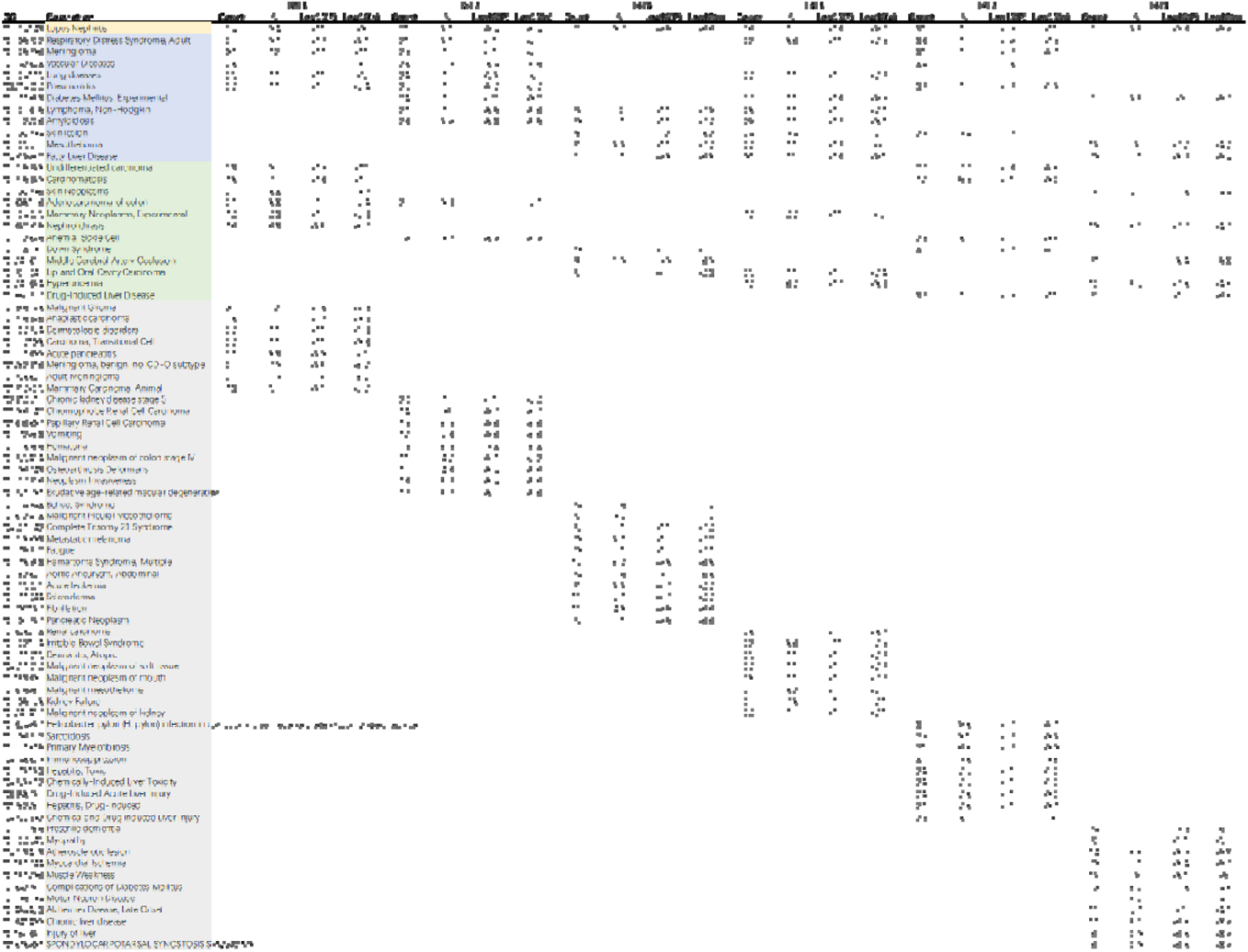
Top 20 clusters with their representative enriched related disease terms before and after homozygous time points

The differential proteins obtained by time point comparison before and after heterozygosity were imported to the Metascape website for enrichment function analysis, as shown in Table 10 below, and all the biological pathways enriched into the top 20 by comparison were listed. We found that there were still many metabolic-related processes, including production of precursor metabolites and energy, alcohol metabolism, degradation of valine, leucine and isoleucine, aldehyde metabolism in cells, lipid catabolism, small molecule metabolism containing nucleobases, carbohydrate metabolism, small molecule catabolism, sulfur-containing compound metabolism, peptide metabolism, organic hydroxyl compound metabolism, amino acid metabolism, and ceramide catabolism. The comparison of the latter two time points with the previous time points enriched the Parkinson’s disease, in addition to some regulatory processes such as oxidative stress, angiogenesis, bone marrow cell differentiation, post-translational protein phosphorylation, epidermal cell differentiation, retinal homeostasis, morphogenesis of sensory organs, and renal development. Subsequently, the differential proteins were also analyzed for gene and disease association by DisGeNET, as shown in Table 11 below. Cancer, transitional cells are enriched in each comparison, already present at a second time point, meningiomas begin to appear at a third time point, and such as fatty liver, senile dementia, motor neuron disease, diarrhea, vascular disease, cancer spread, inflammation, undifferentiated cancer, enzyme disease, pediatric astrocytoma, post-traumatic osteoporosis, hemolytic anemia, fatigue, splenomegaly, hepatomegaly also begin to appear at a second time point. While tumors such as pancreatic tumors, emphysema, chemical and drug-induced liver injury, drug-induced liver disease, toxic hepatitis, chemically induced hepatotoxicity, mesothelioma, adult respiratory distress syndrome, head and neck cancer, colon adenocarcinoma, and pancreatic adenocarcinoma began to appear at three or four time points. Diseases unique to the third time point compared with the previous time point included drug-induced acute liver injury, drug-induced hepatitis, vomiting, tuberculosis-like, idiopathic chronic pancreatitis, Persian Gulf syndrome, Parkinson’s disease, primary hyperthyroidism, atherosclerotic lesions, Guam-type amyotrophic lateral sclerosis, Hartnupp’s disease, dysphagia, hepatocellular carcinoma in children, hepatocellular carcinoma in adults, progressive muscle atrophy, cardiac arrest, and language disorder. Unique diseases that have been enriched since the fourth time point compared to the previous time point include metabolic/homeostatic abnormalities, hepatoblastoma, thinning of the skin, cervical intraepithelial neoplasia, ductal carcinoma of the breast, amnesia, myocarditis, aortic dissection, scleroderma, phenylketonuria, malformed osteoarthropathy, non-Hodgkin’s lymphoma, prostate cancer, leukoencephalopathy, neuropathy, Fanconi anemia, benign meningiomas, precancerous lesions, ductal carcinoma, Down’s syndrome, lupus nephritis, and glaucoma.

**Table 10.**
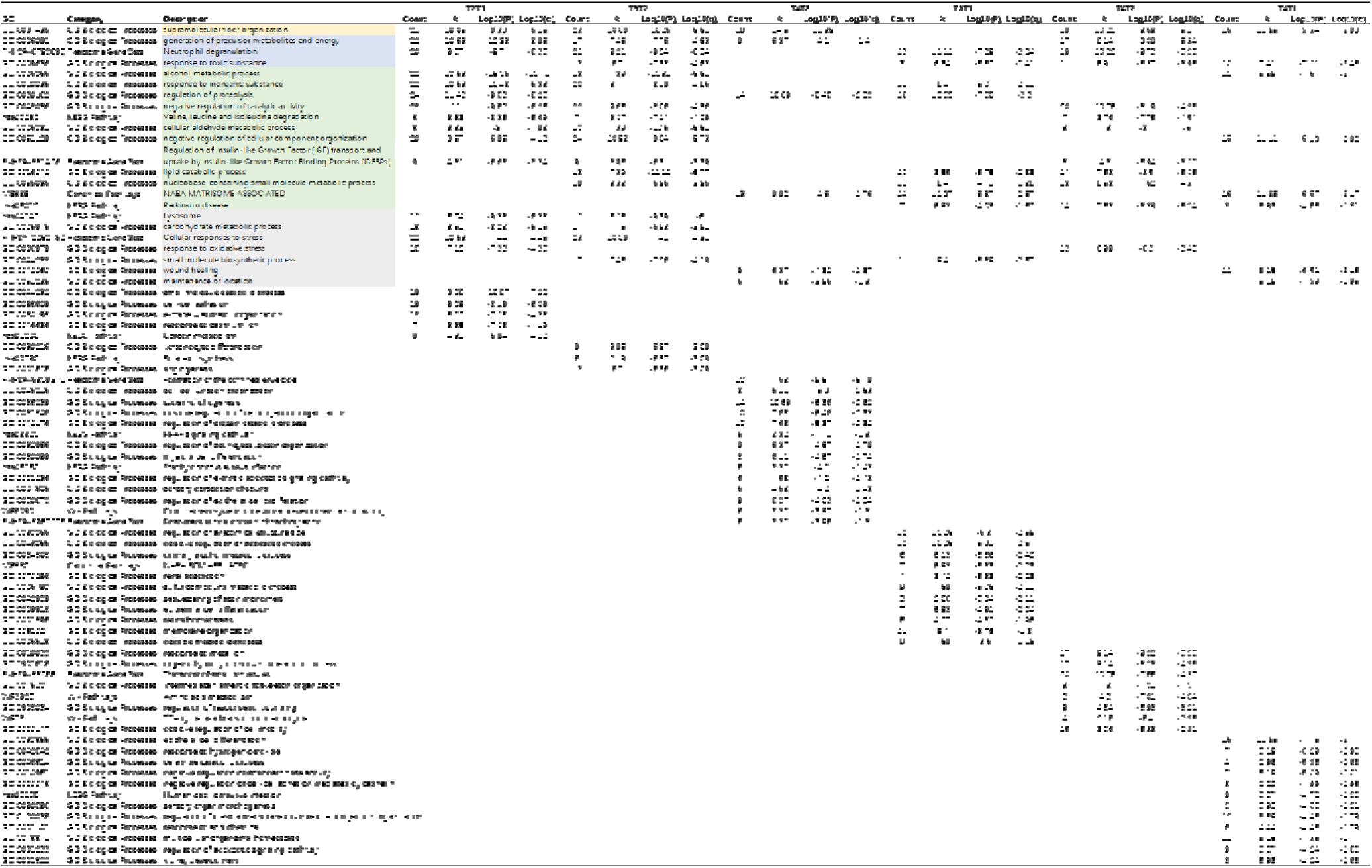
Top 20 clusters with their representative enriched biological processes terms before and after heterozygote time points

**Table 11.**
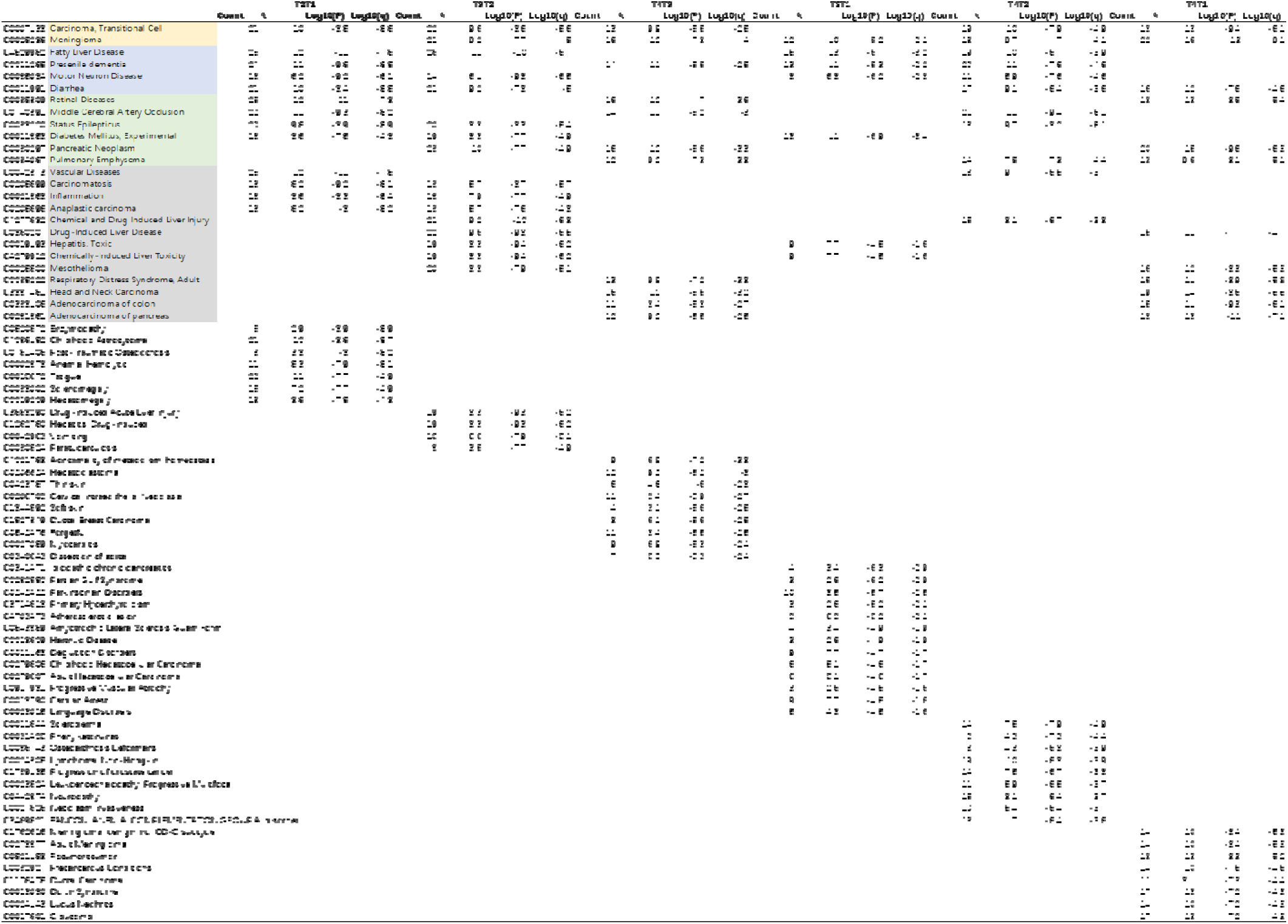
Top 20 clusters with their representative enriched related disease terms before and after heterozygote time points

The intersection of the differential proteins obtained by the homozygote/heterozygote control before and after the treatment is shown in Fig. 14 below. The left figure is the intersection of the differential proteins obtained by the homozygote at each time point, and the right figure is the intersection of the differential proteins obtained by the heterozygote at each time point, and it can be seen that there are many overlapping differential proteins. All genes corresponding to the submitted differential proteins were used as enrichment background for disease or abnormal phenotypes with p values < 0.01, minimum count of 3, and enrichment factor > 1.5 (the enrichment factor being the ratio between the observed count and the accidentally expected count) and grouped into clusters based on their member similarities. The next picture on the left in Figure 15 reflects the enrichment of possible related diseases, including primary myelofibrosis, undifferentiated cancer, hyperuricemia, diabetes, amyloidosis, mesothelioma, fatty liver, skin lesion, Down’s syndrome, meningioma, respiratory distress syndrome, lupus nephritis, sarcomatosis, malignant glioma, vascular disease, pneumonia, cancer spread, undifferentiated cancer, and drug-induced liver disease. On the right side of Fig. 15, tissues and organs to which the differential proteins belong are shown, including colon, bronchus, epithelial cells, testis, germ cells, hepatocytes, adipocytes, small intestine, skin, placenta, tongue, pancreas, adipose tissue, tonsil, breast cells and stomach. The lower left figure in Fig. 15 shows the diseases enriched with differential proteins obtained by comparison before and after time points of heterozygosity, including vascular disease, motor neuron disease, retinal disease, cancer spread, transitional cell carcinoma, chemical and drug-induced liver injury, chemical hepatotoxicity, drug-induced acute liver injury, drug-induced hepatitis, drug-induced liver disease, toxic hepatitis, persistent epileptic state, fatty liver, Alzheimer’s disease, meningioma, pancreatic cancer, middle cerebral artery occlusion, respiratory distress syndrome, colon adenocarcinoma, and pancreatic tumor. The bottom right panel of Fig. 15 shows the tissues and organs to which the differential proteins belong, including liver, colon, bronchial epithelial cells, testicular germ cells, hepatocytes, adipocytes, small intestine, skin, placenta, tongue, pancreas, adipose tissue, tonsil, mammary gland cells and stomach.

**Fig. 14.**
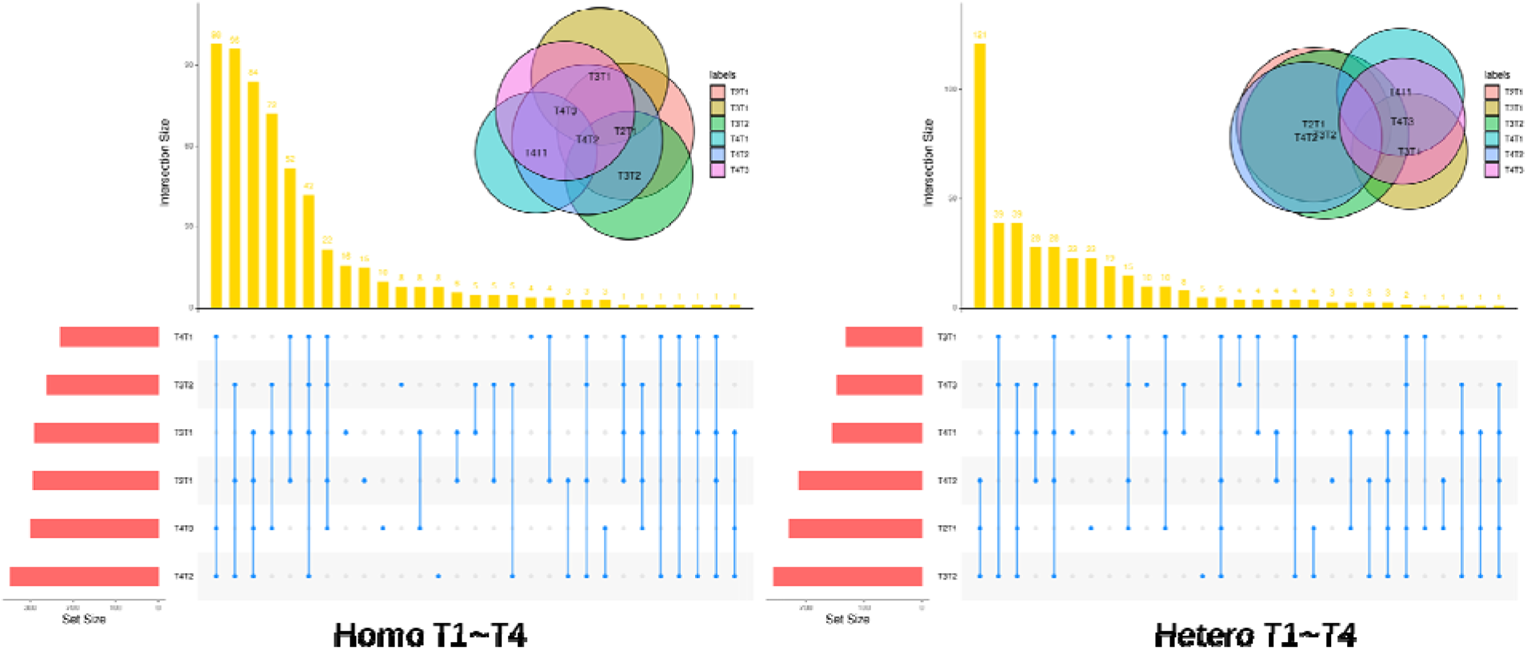
The interaction between differential proteins of homozygotes/heterozygotes before and after different time points

**Fig. 15.**
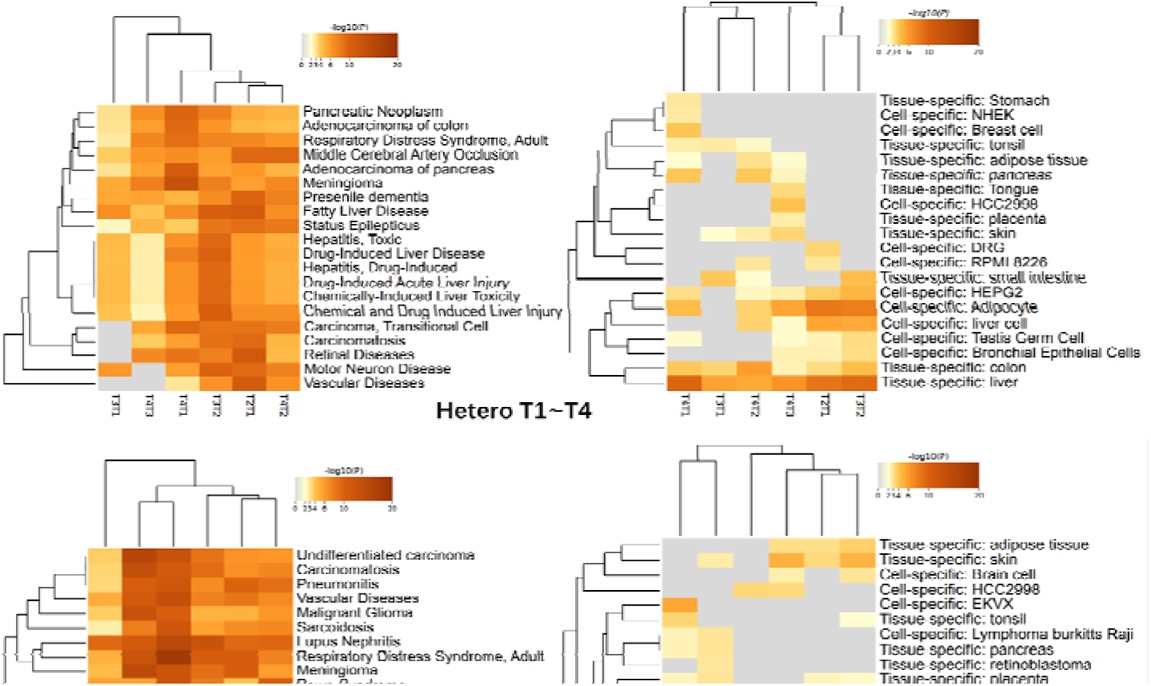
Summary of enrichment analysis in DisGeNET、PaGenBase(Homozygotes/heterozygotes)

### Self-control analysis of three p53 gene knockout mice

We selected three mice numbered 28, 32 and 33 and involved three early time points, including 22 days T1, 42 days T2 and 62 days T3 after weaning, to compare and analyze the single self-control before and after, and screen out the respective differential proteins according to the previous criteria, and then conduct the related possible disease analysis through the String link https://monarchinitiative.org/. It was found that the pathways for each mouse to accumulate the related diseases at each time point were not completely the same, and there were some differences, as shown in Table 12 below.

**Table 12.**
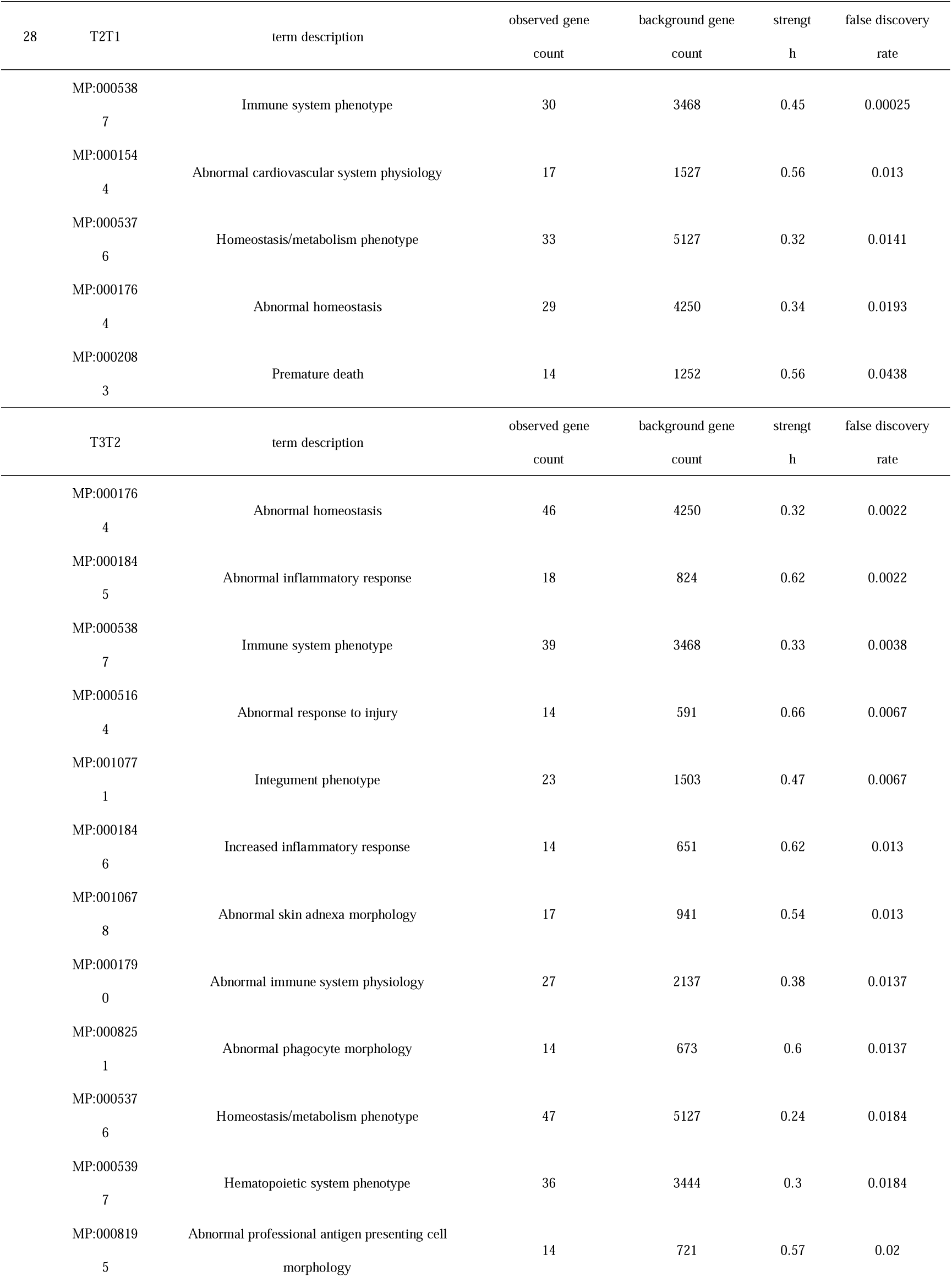
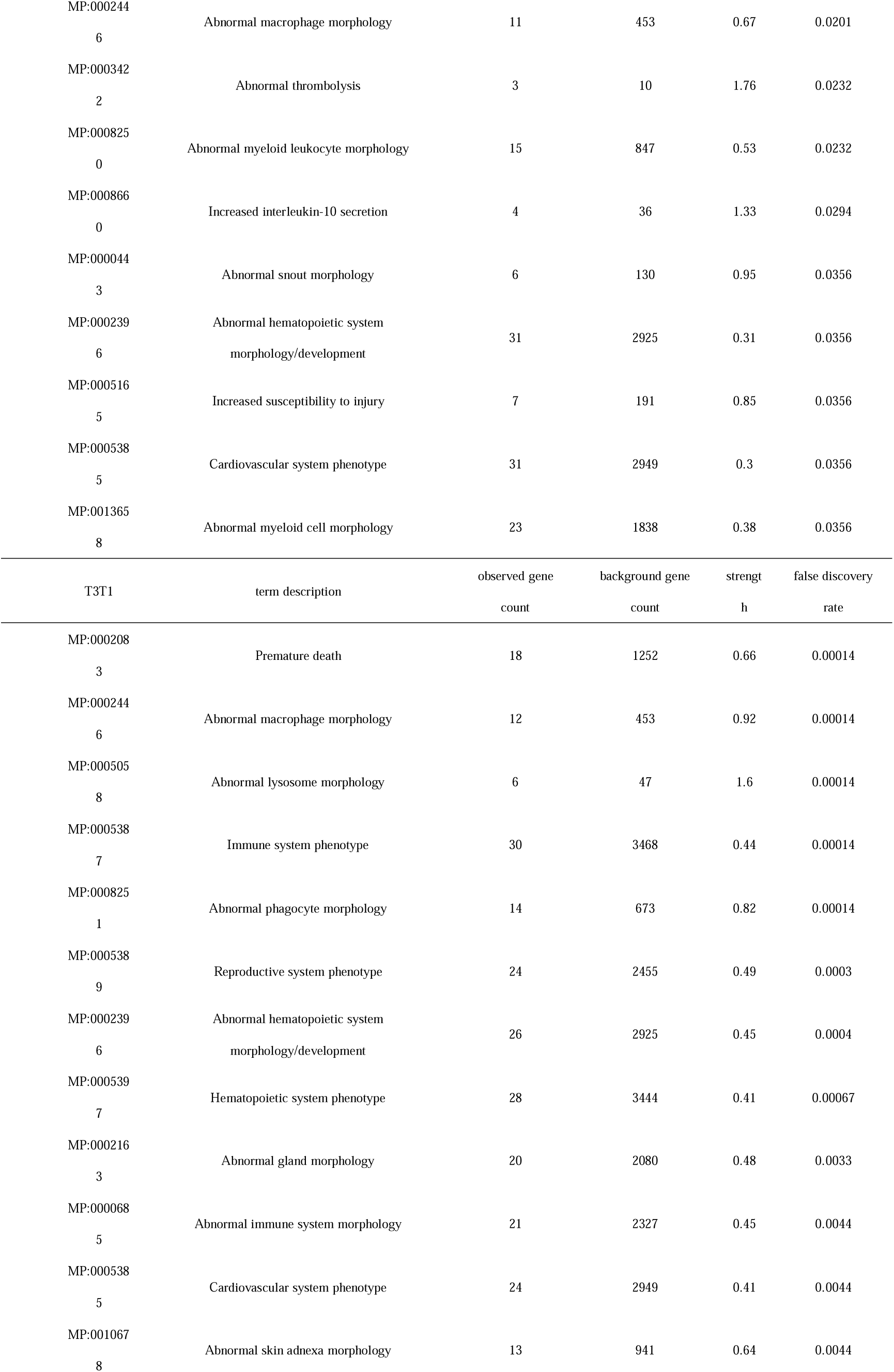
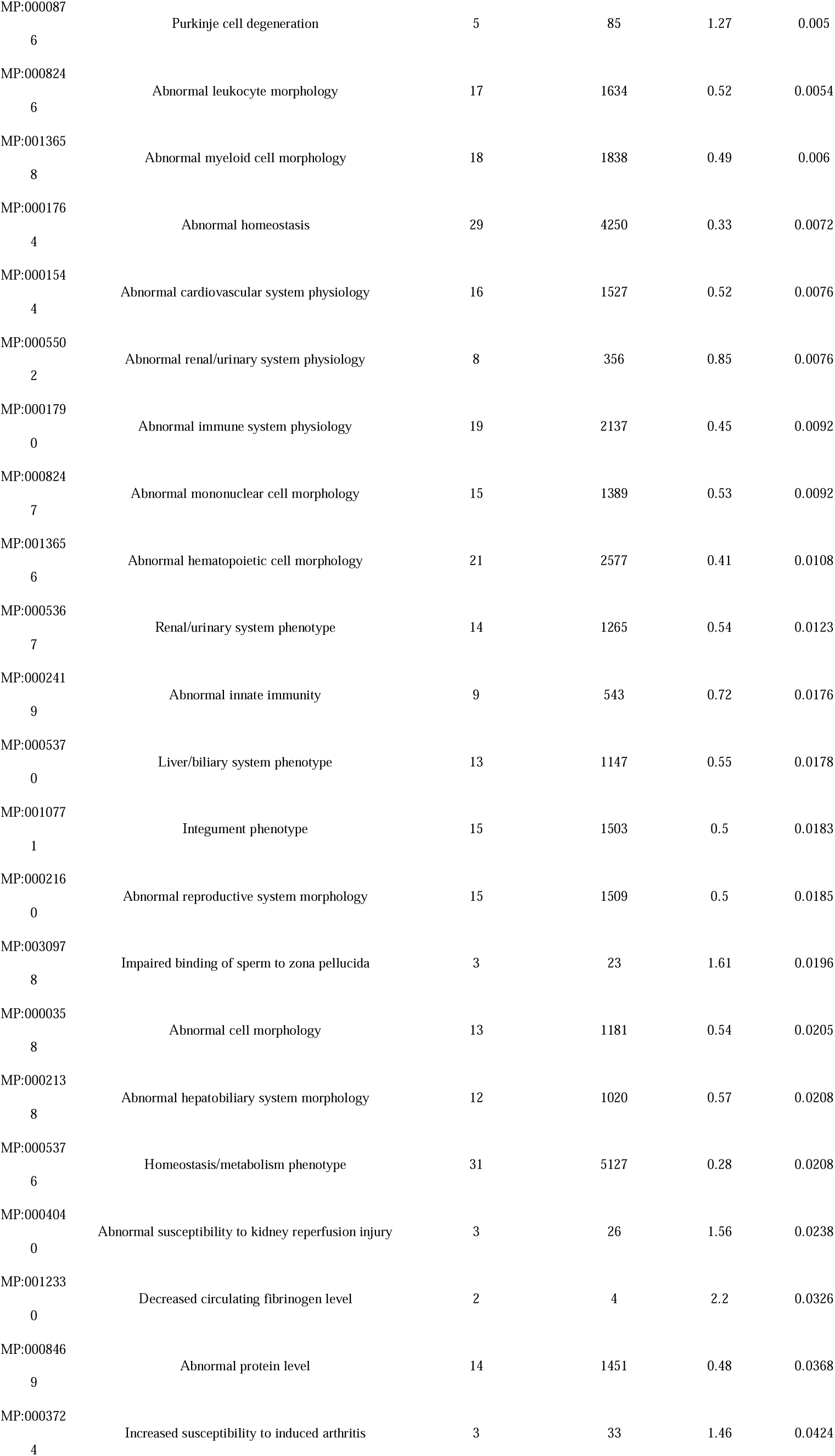
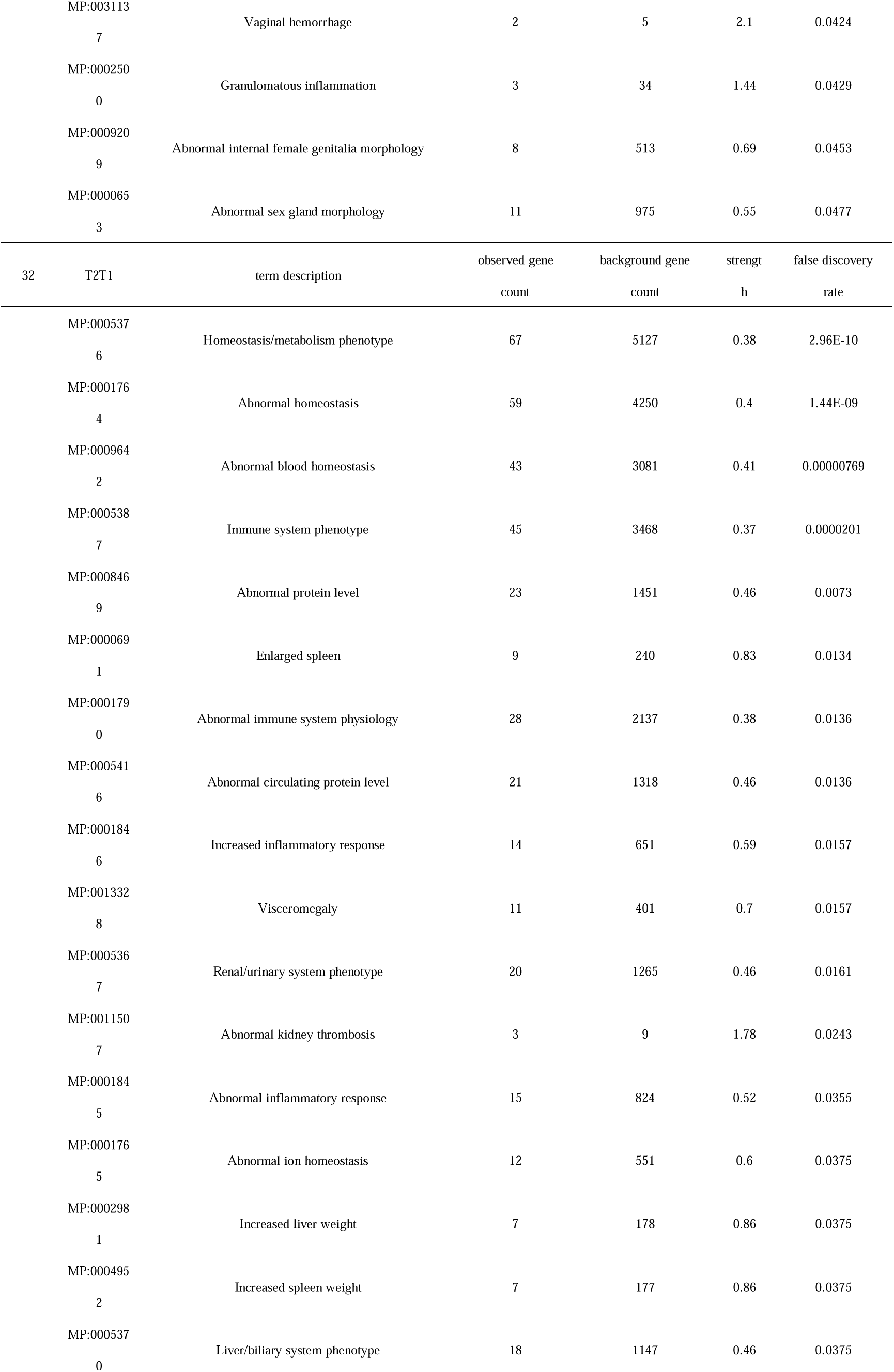
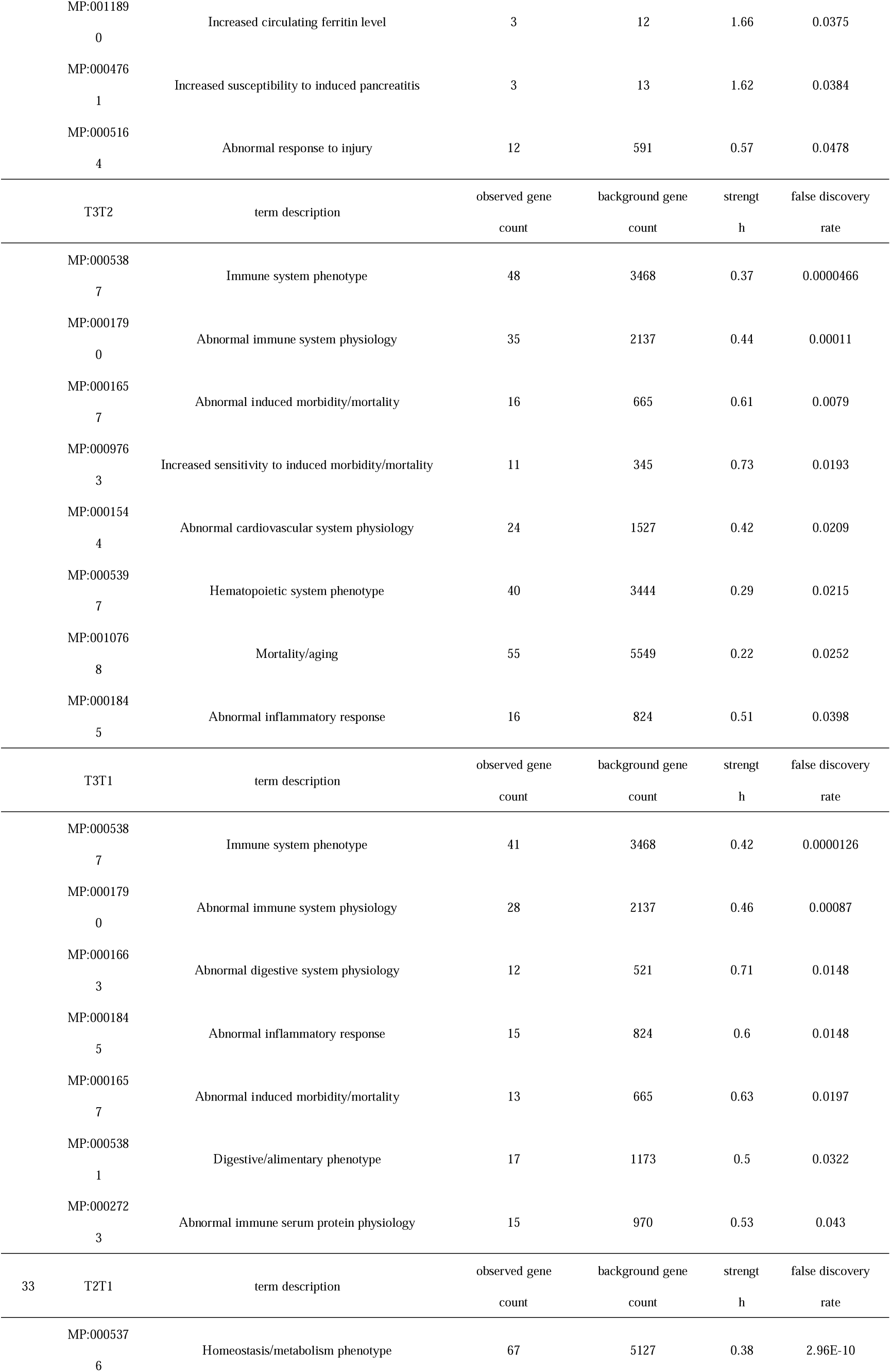
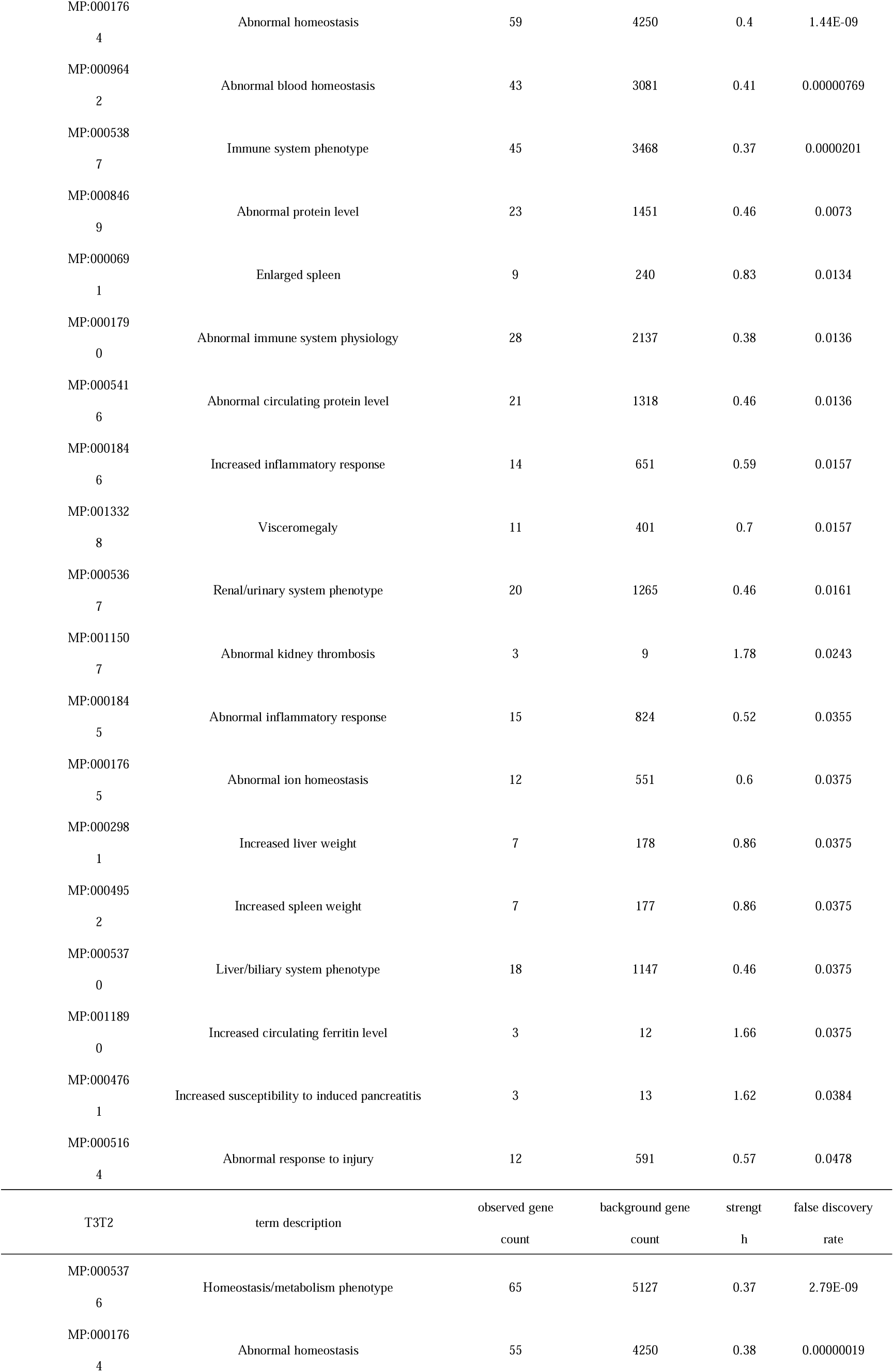
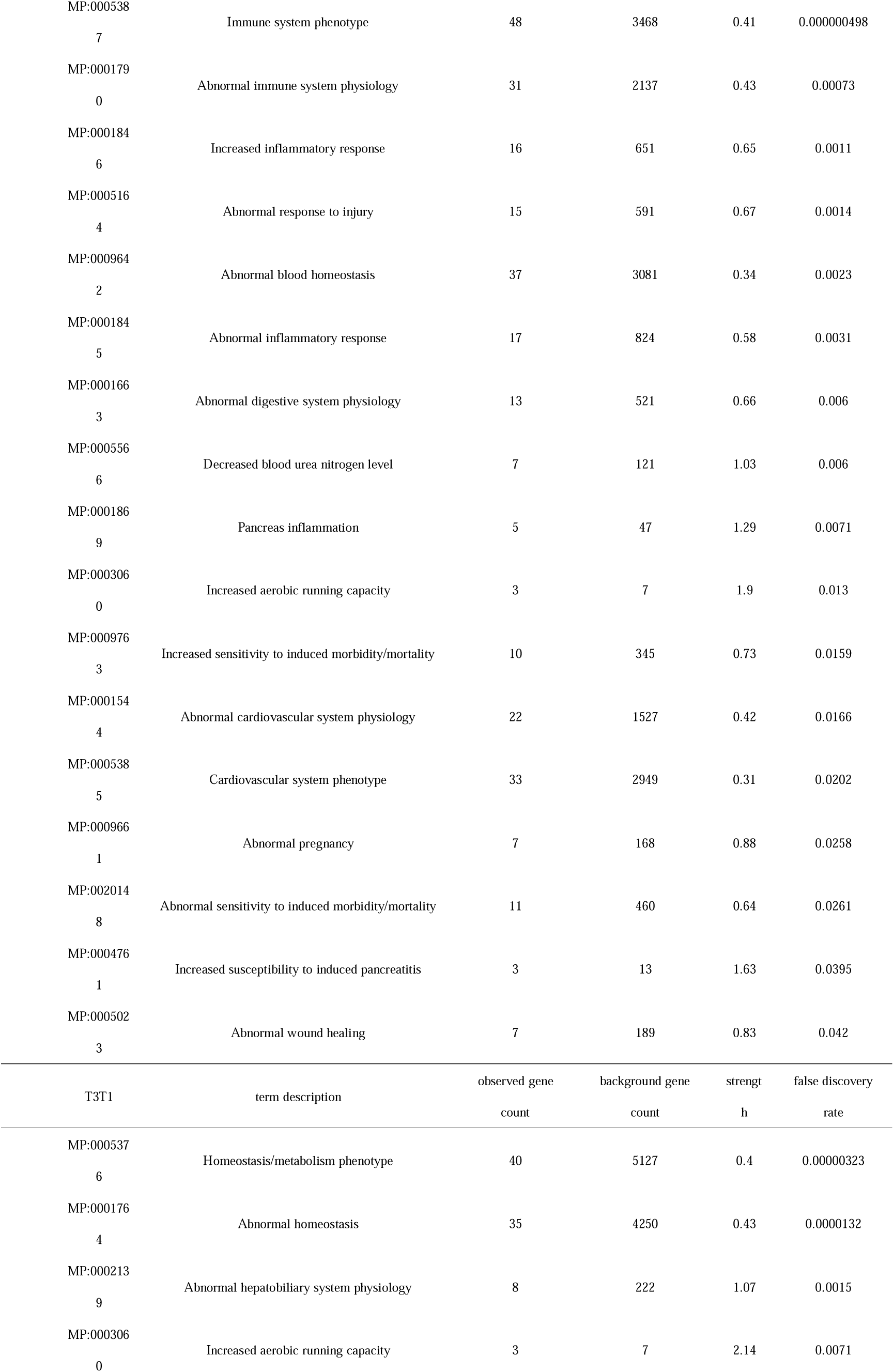
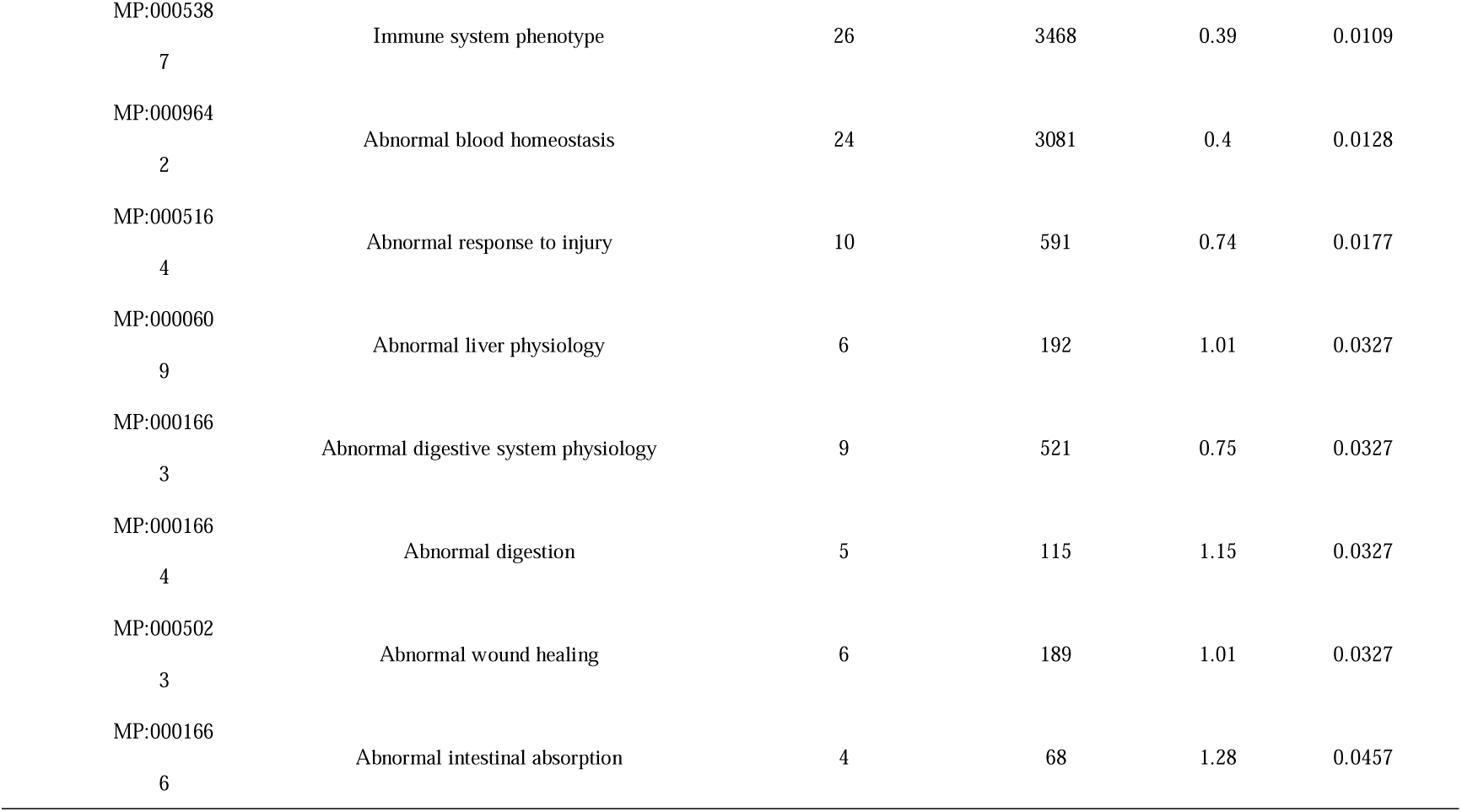
Differential protein-related diseases analysis of mice self-comparison (Monarch)

## Discussion

In this experiment, we imported the obtained differential proteins into STRING website for ontology analysis of mammalian phenotype (The Mammalian Phenotype Ontology, Monarch). This function relies on the Monarch website. The Monarch Initiative is a platform for data integration and analysis, connecting phenotypes with cross-species genotypes, and connecting basic and applied research with semantic-based analysis. Exploring the correlation between phenotypic outcomes, disease and genetic variation, as well as environmental factors, the platform creates a biological ontology based on this, which together enables computational analysis of complex and semantic integration of gene, genotype, variant, disease and phenotypic data, identification of animal models of human disease by phenotypic similarity, phenotypically driven computational support for differential diagnosis, and transformation research. At the same time, we introduced the differential proteins obtained from normal growth and development rats in the previous published articles. Monarch is not enriched for related diseases, which may also explain the abnormal effects on mouse body after p53 gene knockout, and it may have a tendency to cause diseases. The urine protein group can reflect the stable state disorder and imbalance of the mouse body after gene knockout, and there are even signs of various diseases, which may be the reason for its multiple tumors, different phenotypes and final premature death.

